# Learning structural heterogeneity from cryo-electron sub-tomograms with tomoDRGN

**DOI:** 10.1101/2023.05.31.542975

**Authors:** Barrett M. Powell, Joseph H. Davis

**Affiliations:** Department of Biology; Program in Computational and Systems Biology Massachusetts Institute of Technology Cambridge, MA 02139

## Abstract

Cryo-electron tomography (cryo-ET) allows one to observe macromolecular complexes in their native, spatially contextualized environment. Tools to visualize such complexes at nanometer resolution via iterative alignment and averaging are well-developed but rely on assumptions of structural homogeneity among the complexes under consideration. Recently developed downstream analysis tools allow for some assessment of macromolecular diversity but have limited capacity to represent highly heterogeneous macromolecules, including those undergoing continuous conformational changes. Here, we extend the highly expressive cryoDRGN deep learning architecture, originally created for cryo-electron microscopy single particle analysis, to sub-tomograms. Our new tool, tomoDRGN, learns a continuous low-dimensional representation of structural heterogeneity in cryo-ET datasets while also learning to reconstruct a large, heterogeneous ensemble of structures supported by the underlying data. Using simulated and experimental data, we describe and benchmark architectural choices within tomoDRGN that are uniquely necessitated and enabled by cryo-ET data. We additionally illustrate tomoDRGN’s efficacy in analyzing an exemplar dataset, using it to reveal extensive structural heterogeneity among ribosomes imaged *in situ*.

## INTRODUCTION

Life relies on an array of large, dynamic macromolecular complexes to carry out essential cellular functions. The conformational flexibility and compositional variability in these complexes allow cells to mount targeted molecular responses to various stresses and stimuli. Structural biology has long aimed to visualize these diverse structures with the goals of gaining mechanistic insights into these responses and testing hypotheses related to macromolecular structure-function relationships. In pursuit of this goal, cryo-electron microscopy (cryo-EM) has proven to be a powerful tool for visualizing purified complexes with high resolution (Bai *et al*., 2015; Murata and Wolf, 2018). In cryo-EM, ∼10^4^-10^7^ individual particles are imaged by transmission electron microscopy (TEM), each from a single unknown projection angle. Single particle analysis (SPA) is then used to simultaneously estimate the most likely projection angle for each particle image and the *k*≥1 distinct 3-D volumes of the target complex, which, when projected to 2-D, are most likely to have produced the source dataset (Cheng *et al*., 2015). More recently, a number of tools have leveraged SPA datasets to deeply explore structural heterogeneity within these complexes (Chen and Ludtke, 2021; Dashti *et al*., 2020; Kinman *et al*., 2023; Punjani and Fleet, 2021; Sun *et al*., 2022; Zhong *et al*., 2021), dramatically expanding the range of insights and testable biological hypotheses that can be derived from cryo-EM.

Cryo-electron tomography (cryo-ET) is a related imaging modality wherein a sample is repeatedly imaged from several known projection angles, enabling the reconstruction of a 3-D tomogram (Asano *et al*., 2016). As such, cryo-ET disentangles particles that overlap along a projection axis and enables the nanometer-scale 3-D visualization of highly complex samples, including subcellular volumes. Thus, cryo-ET affords the opportunity to inspect macromolecular structures in their native cellular context (Gemmer *et al*., 2023; Hoffmann *et al*., 2022; Lovatt *et al*., 2022; Xue *et al*., 2022), in contrast with cryo-EM’s typical requirement that particles be isolated from cells and purified.

Sub-tomogram averaging (STA), a particle averaging approach analogous to SPA, is often employed in cryo-ET data processing. In STA, individual 3-D volumes, each a sub-tomogram corresponding to a unique particle, are extracted from the back-projected tilt series and are iteratively aligned to produce an average particle volume with increased SNR and resolution (Bharat and Scheres, 2016; Castano-Diez and Zanetti, 2019; Pyle and Zanetti, 2021; Zhang, 2019). Critically, STA can therefore offer insights to native protein complexes, enabling hypothesis generation in identifying unknown associated factors or novel complex ultrastructure.

As with SPA, several tools have recently been developed to characterize heterogeneity among individual particles relative to the global average (Castano-Diez *et al*., 2012; Harastani *et al*., 2021; 2022; Himes and Zhang, 2018; Stolken *et al*., 2011), either during or after STA. Although these approaches have proven fruitful in answering specific biological questions such as nucleosome flexibility (Harastani *et al*., 2021; 2022), and ribosome heterogeneity (Himes and Zhang, 2018; Xue *et al*., 2022), each approach has specific constraints that limit their generality. For example, sub-tomogram PCA (Himes and Zhang, 2018) assumes heterogeneity can be modeled as a linear combination of voxel intensity, normal mode analysis (Harastani *et al*., 2021) requires *a priori* knowledge of an atomic model or density map to compute normal modes, and optical flow (Harastani *et al*., 2022) is inherently limited to conformational changes of the target particle in which the total voxel intensity across each sub-tomogram remains approximately constant. An unbiased and expressive tool to analyze heterogeneity is therefore highly desirable, particularly for *in situ* discovery of unexpected cofactors whose identity, binding site, and occupancy may be unknown.

Here, we introduce tomoDRGN (<u>D</u>eep <u>R</u>econstructing <u>G</u>enerative <u>N</u>etworks), a deep learning framework designed to learn a continuously generative model of per-particle conformational and compositional heterogeneity from cryo-ET datasets. TomoDRGN is related to our well-characterized cryoDRGN software (Kinman *et al*., 2023; Zhong *et al*., 2021), and therefore shares many overall design, processing, and analysis philosophies. As input, tomoDRGN uses particle images and corresponding metadata from upstream STA tools (**Fig. 1a**). It then learns to simultaneously embed each particle within a continuous low dimensional latent space and to reconstruct the corresponding unique 3-D volume (**Fig. 1b**). We have additionally developed and integrated software tools to visualize and interpret these outputs, and to integrate tomoDRGN outputs with external processing software for subsequent analyses, including contextualizing the tomoDRGN generated volumes within the *in situ* cellular tomography data.

**Figure 1:**
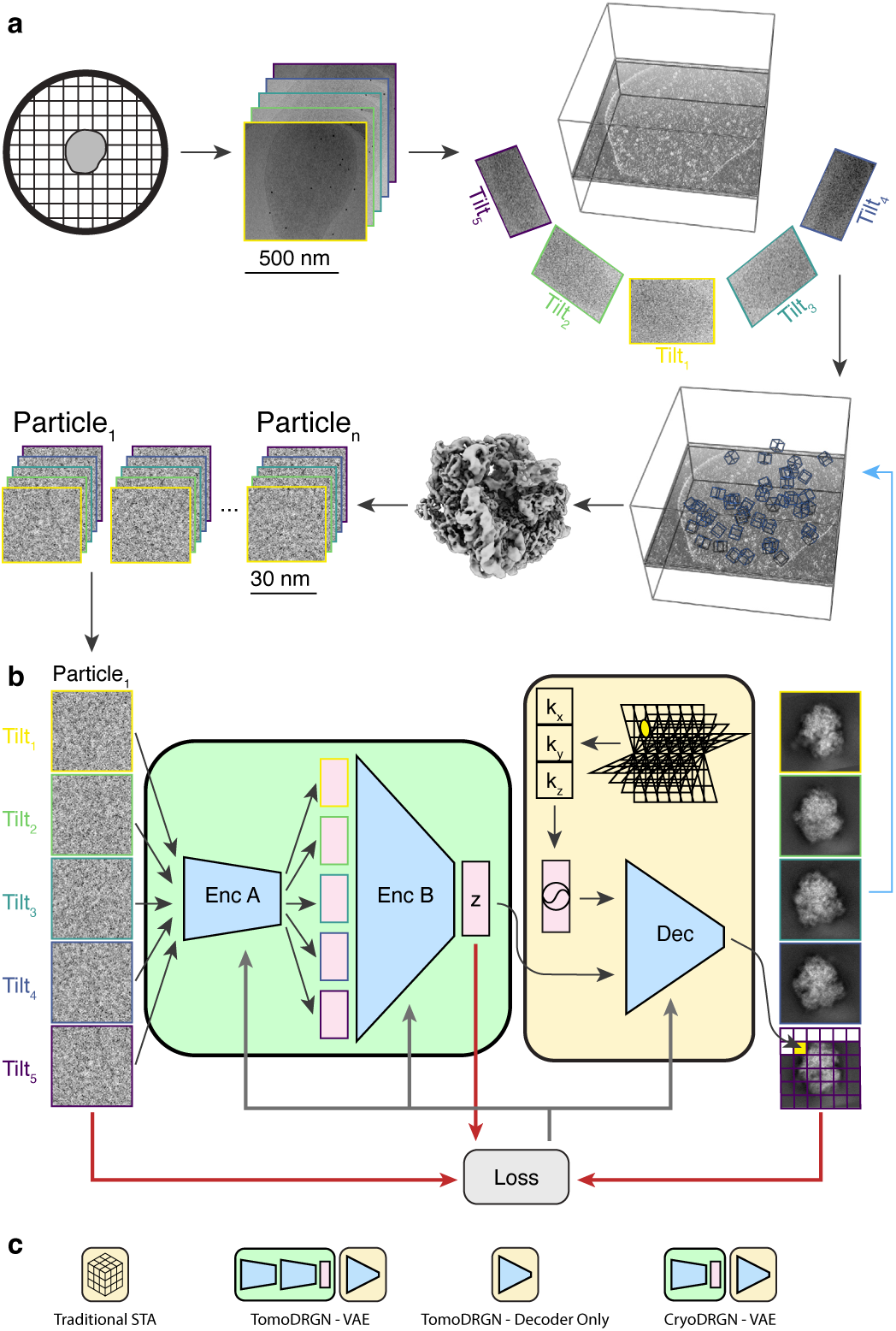
A neural network architecture to analyze structurally heterogeneous particles imaged by cryo-ET. (a) A typical sample and data processing workflow to produce tomoDRGN inputs. The sample (*e.g.*, a bacterial cell) is applied to a grid, plunge frozen, and optionally thinned. A series of TEM images of a target region are collected at different stage tilt angles. A tomographic volume is reconstructed using weighted back-projection of all tilt images. Instances of the target particle are identified (blue boxes) and extracted as 3-D voxel arrays. Iterative sub-tomogram averaging (STA) is used to reconstruct a consensus density map. Per-particle 2-D tilt images are then re-extracted from the source tilt series images and parameters (*e.g.* pose, defocus, etc.) estimated from STA are associated with the images. **(b)** The tomoDRGN network architecture and training design. Each particle’s set of tilt images are independently passed through encoder A (Enc A), then jointly passed through encoder B (Enc B), thereby mapping all tilt images of a particle to one embedding (*z*) in a low dimensionality latent space. The decoder network (Dec) uses the latent embedding and a featurized voxel coordinate to decode a corresponding set of images pixel-by-pixel. Note that the decoder can learn a homogeneous structure by excluding the encoder module. The network is trained using a loss function (grey arrows) that depends on the input images, reconstructed images, and *z* (red arrows). **(c)** Graphical signposts for volumes generated or analyzed by distinct reconstruction tools. These signposts are used throughout this manuscript when volumes are displayed to clarify how they were generated.

## RESULTS

### A deep learning framework to reconstruct heterogeneous volumes from TEM tilt-series data

TomoDRGN was designed to efficiently train a neural network capable of: 1) embedding a collection of particles (each represented by multiple TEM images collected at different stage tilts) in a learned, continuous, low-dimensional latent space informed by structural heterogeneity; and 2) generating a 3-D volume for each individual particle using these embeddings. By design, cryoDRGN is unsuited for this task as it maps individual images to unique latent embeddings, which is expected for cryo-EM single particle datasets. Thus, cryoDRGN is not constrained to map multiple tilt images of the same particle to consistent regions of latent space, leading to uninterpretable learned latent spaces and generated volumes (see Discussion).

To handle tilt-series data, we employed a variational autoencoder (VAE) framework (Kingma and Welling, 2013), featuring a purpose-built two-part encoder network feeding into a coordinate-based decoder network (Bepler *et al*., 2019; Zhong *et al*., 2019) (**Fig. 1b**). For each particle, the encoder network first uses encoder A (per tilt image) as a “feature extractor” to generate a unique intermediate embedding for each tilt image in a manner directly analogous to cryoDRGN’s encoder network. Encoder B then integrates these intermediate embeddings into a single latent embedding for the particle. The decoder network is supplied with this integrated latent embedding and a featurized voxel coordinate to reconstruct the signal at that coordinate. As in cryoDRGN, these operations are performed in reciprocal space. With this design, repeatedly evaluating the decoder network at multiple coordinates should allow for a rasterized reconstruction of the set of tilt images originally supplied to the encoder. Following a standard VAE (Kingma and Welling, 2013), we designed the network to be trained by minimizing a reconstruction loss between input and reconstructed images, and a latent loss quantified by the KL-divergence of the latent embedding from a standard normal distribution.

Once trained, we expected a tomoDRGN network to enable detailed and systematic interrogation of structural heterogeneity within the input dataset. For example, similar to cryoDRGN, tomoDRGN’s learned latent space could be visualized either directly along any sets of latent dimensions or using a dimensionality reduction technique such as UMAP (Becht *et al*., 2018), where we have empirically found that distinct clusters often correspond to compositionally heterogeneous states, and diffuse, unfeatured distributions correspond to continuous structural variation. We reasoned that latent embeddings, sampled individually or following a well-populated path in latent space, could then be passed to the decoder to generate corresponding 3-D volumes for direct visualization. We predicted additional analysis could be performed in 3-D voxel space using standard cryoDRGN tools (Kinman *et al*., 2023; Sun *et al*., 2022). To complement tomoDRGN, we also constructed interactive tools to visualize and analyze heterogeneity in the spatial context of the original tomograms. Finally, we built tools to isolate particle subsets of interest for subsequent refinement with traditional STA software (**Fig. 1c**) as an iterative approach we speculated could maximize the value of a tomographic dataset.

### Sub-tomogram-specific image processing approaches

Having conceived the general tomoDRGN framework, we next considered additional image processing procedures that we hypothesized might improve model quality and computational performance. First, we noted that STA software tools commonly implement weighting schemes to model the signal-to-noise ratio (SNR) of each image as a function of the image tilt angle (*i.e.*, electron pathlength through the sample) and cumulative dose (*i.e.*, accumulated radiative damage) (Bharat *et al*., 2015; Grant and Grigorieff, 2015; Tegunov *et al*., 2021). Thus, we followed standard formulations for tilt weighting as the cosine of the stage tilt angle and dose weighting using fixed exposure curves, and we incorporated such weights into the reconstruction error calculated in tomoDRGN’s decoder network (**Extended Data Fig. 1a**). We expected such an approach would effectively downweigh the reconstruction loss of highly tilted and radiation damaged images, particularly at high spatial frequencies (**Extended Data Fig. 1b-d**).

Second, tomoDRGN’s coordinate-based decoder is trained by evaluating a set of spatial frequencies per tilt image that, by default, is identical for all tilt images (*i.e.*, independent of cumulative dose imparted at each tilt). However, prior work has shown that the SNR at a given spatial frequency can be maximized at an optimal electron dose (Hayward and Glaeser, 1979) and that during cryo-EM movie alignment, filtering spatial frequencies in each frame by their optimal dose can improve the aligned micrograph quality (Glaeser, 1979; Grant and Grigorieff, 2015). We therefore implemented a scheme applying optimal dose filtering to Fourier coordinates evaluated by the decoder during model training (**Extended Data Fig. 1a**). We expected that such filtering would restrict the set of spatial frequencies evaluated during decoder training without sacrificing 3-D reconstruction accuracy, thereby decreasing the computational burden of model training, particularly for high resolution datasets at large box sizes (**Extended Data Fig. 1b-d**).

Finally, real-world datasets frequently contain particles missing some tilt images, often due to upstream micrograph filtering (**Extended Data Fig. 2a**). To flexibly handle such nonuniform input data, we implemented an approach that surveys the dataset for the fewest tilt images associated with a single particle (*n*), then randomly samples *n* tilt images from each particle during model training and evaluation (**Extended Data Fig. 2b**). Because this approach subsets and permutes tilt images at random, encoder B must learn a permutation-invariant function mapping from encoder A’s output (per tilt image) to the final latent space (per particle), and we hypothesized that this permutation-invariant learning goal might provide added regularization that could decrease overfitting.

### TomoDRGN robustly recovers simulated heterogeneity

To judge the efficacy of these architectural choices, we simulated (Baxter *et al*., 2009) cryo-ET particle stacks (see Methods) corresponding to four assembly states (B-E) of the ribosomal large subunit (LSU) from *E. coli* (Davis *et al*., 2016; Davis and Williamson, 2017) (**Fig. 2a**). We initially tested the ability of the isolated decoder network to perform a homogeneous reconstruction of the class E particles (*i.e.*, no encoder was trained, and no latent space learned). We observed rapid convergence of the decoder network, with it reproducing the ground-truth density maps within 10 epochs (**Fig. 2b**).

**Figure 2:**
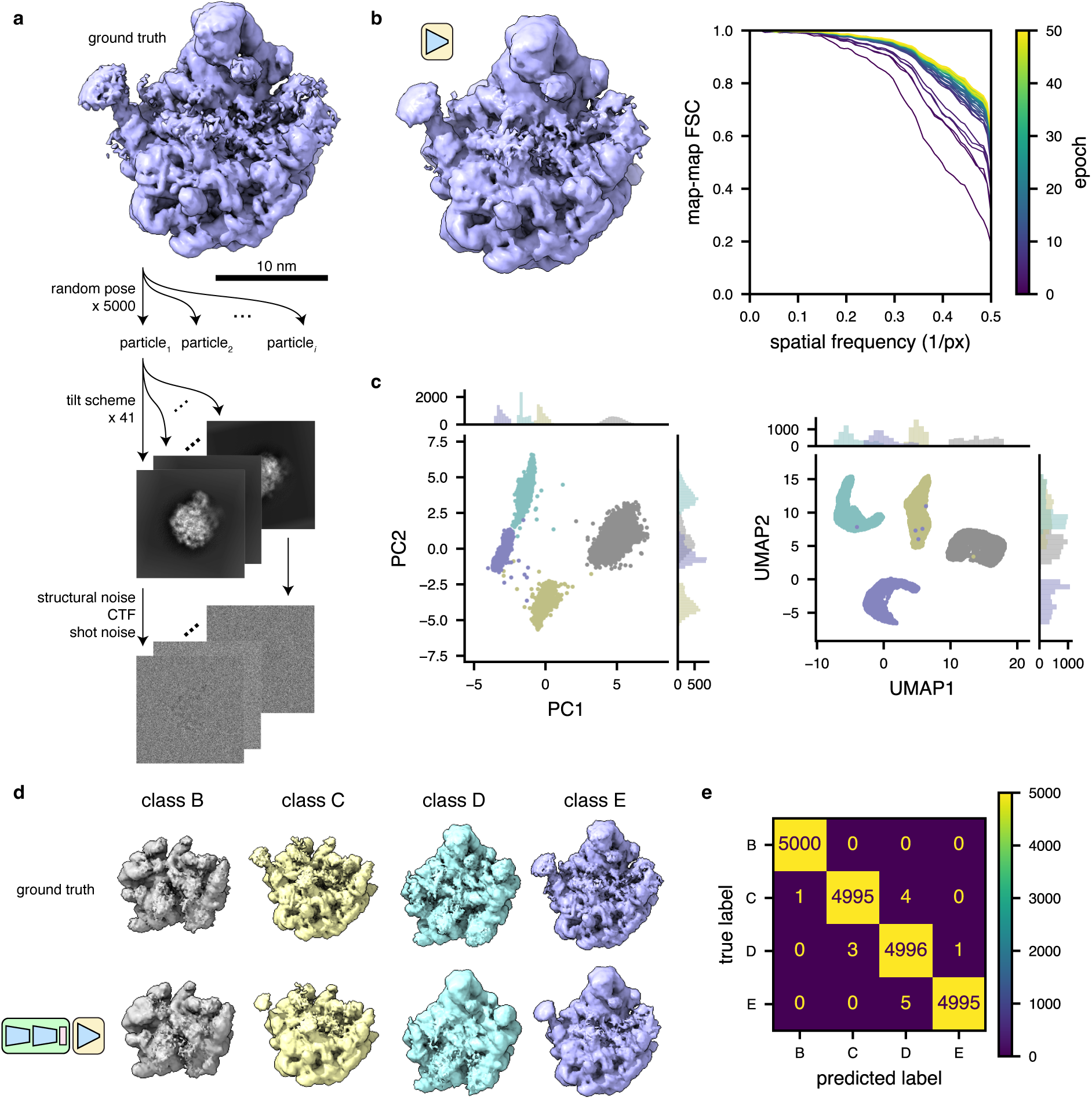
TomoDRGN recovers known heterogeneity in simulated datasets. **(a)** Illustration of the method used to simulate tilt series particle stacks corresponding to four assembly states (B-E) (Davis *et al*., 2016) of the bacterial large ribosomal subunit. **(b)** Left, a tomoDRGN homogeneous network reconstruction of the simulated class E dataset after 50 epochs of training at a resolution of 3.55 Å/px. Right, FSC between the tomoDRGN reconstruction and the ground truth volume at each of 50 epochs of training (purple to yellow). **(c)** First two principal components (left) and UMAP embeddings (right) of tomoDRGN latent space when trained on the simulated four class dataset, colored by *k*=4 *k*-means classification of latent space. **(d)** Ground truth ribosomal volumes (top) and corresponding tomoDRGN-reconstructed volumes (bottom) sampled from the median latent encoding of each of the *k*=4 *k*-means classes in (c). **(e)** Confusion matrix of *k*-means clustering class labels from (c) against ground truth class labels.

To assess tomoDRGN’s ability to faithfully embed and reconstruct structurally heterogeneous 3-D volumes, we next trained the full VAE network using particle stacks containing a labeled mixture of all four LSU structural classes. After training for 24 epochs, we observed four distinct clusters of latent embeddings by PCA and UMAP (**Fig. 2c**). Furthermore, the decoder network generated volumes from the center of each latent cluster that were consistent with the ground truth volumes (**Fig. 2d**). Finally, we quantified the fidelity of the embeddings to their corresponding ground truth volume classes on a per-particle basis. We observed a nearly one-to-one mapping between tomoDRGN particle embeddings and the correct ground truth class (**Fig. 2e**), indicating that the tomoDRGN network effectively learned discrete structural heterogeneity without supervision.

We next assessed the benefits of our aforementioned reconstruction loss weighting, lattice coordinate filtering, and random tilt sampling approaches. Testing the weighting and filtering schemes on the homogeneous reconstruction of the LSU class E ribosomes, we observed modest improvements to final resolution with either or both schemes over using neither. Notably, however, the lattice coordinate filtering scheme led to large reductions in wall clock runtime and GPU memory utilization (**Extended Data Fig. 1c-e**, **Supplementary Table 1**). To assess the efficacy of the random sampling scheme, we compared heterogeneous networks trained on the 4-class LSU dataset with and without random tilt sampling. We observed higher average volume correlation coefficients (CC) for tomoDRGN volumes against ground truth volumes when using random sampling. Random sampling also provided our hypothesized robustness to model overfitting compared to sequential tilt sampling, as evidenced by the stable and elevated average CCs during further model training (**Extended Data Fig. 2c**). Finally, using the random sampling scheme, we observed an interpretable and well-featured latent space, even when using as few as 11 of the 41 available tilt images for each particle (**Extended Data Fig. 2d-e**). We additionally measured the accuracy and consistency of volumes generated from each such latent embedding to the corresponding ground truth volume, per particle per epoch, again observing robust performance with the random sampling scheme (**Extended Data Fig. 2f**). Notably, each of these metrics exhibited a dramatic drop in quality when only using a single tilt sampled per particle. This observation was consistent with our prediction that the mapping of one image to one latent embedding would be unsuitable for tilt series data.

Combined, these strategies allowed efficient and flexible analysis of diverse input datasets, and we have benchmarked tomoDRGN performance for a range of network architectures (**Extended Data Figs. 3-4**, **Supplemental Tables 2-4**). Generally, we observe that performance is robust to network architecture hyperparameters, with slight improvements for deeper and narrower encoder A modules, and wider and shallower decoder modules.

### TomoDRGN uncovers structurally heterogeneous ribosomes imaged *in situ*

We next assessed tomoDRGN’s performance on the publicly available cryo-ET dataset EMPIAR-10499 (Tegunov *et al*., 2021), using it to analyze heterogeneity among chloramphenicol-treated ribosomes imaged in the bacterium *Mycoplasma pneumoniae*. Following published STA methods (Tegunov *et al*., 2021), we reproduced a Nyquist-limited ∼3.5 Å resolution reconstruction of the 70S ribosome (**Fig. 3a**). We subsequently extracted corresponding ribosome images from the aligned tilt micrographs and used this particle stack to train a homogeneous tomoDRGN model. The tomoDRGN-reconstructed volume recapitulated high-resolution features observed in the STA map (**Fig. 3a-c**), highlighting the tomoDRGN decoder network’s ability to learn to accurately represent high-resolution structures.

**Figure 3:**
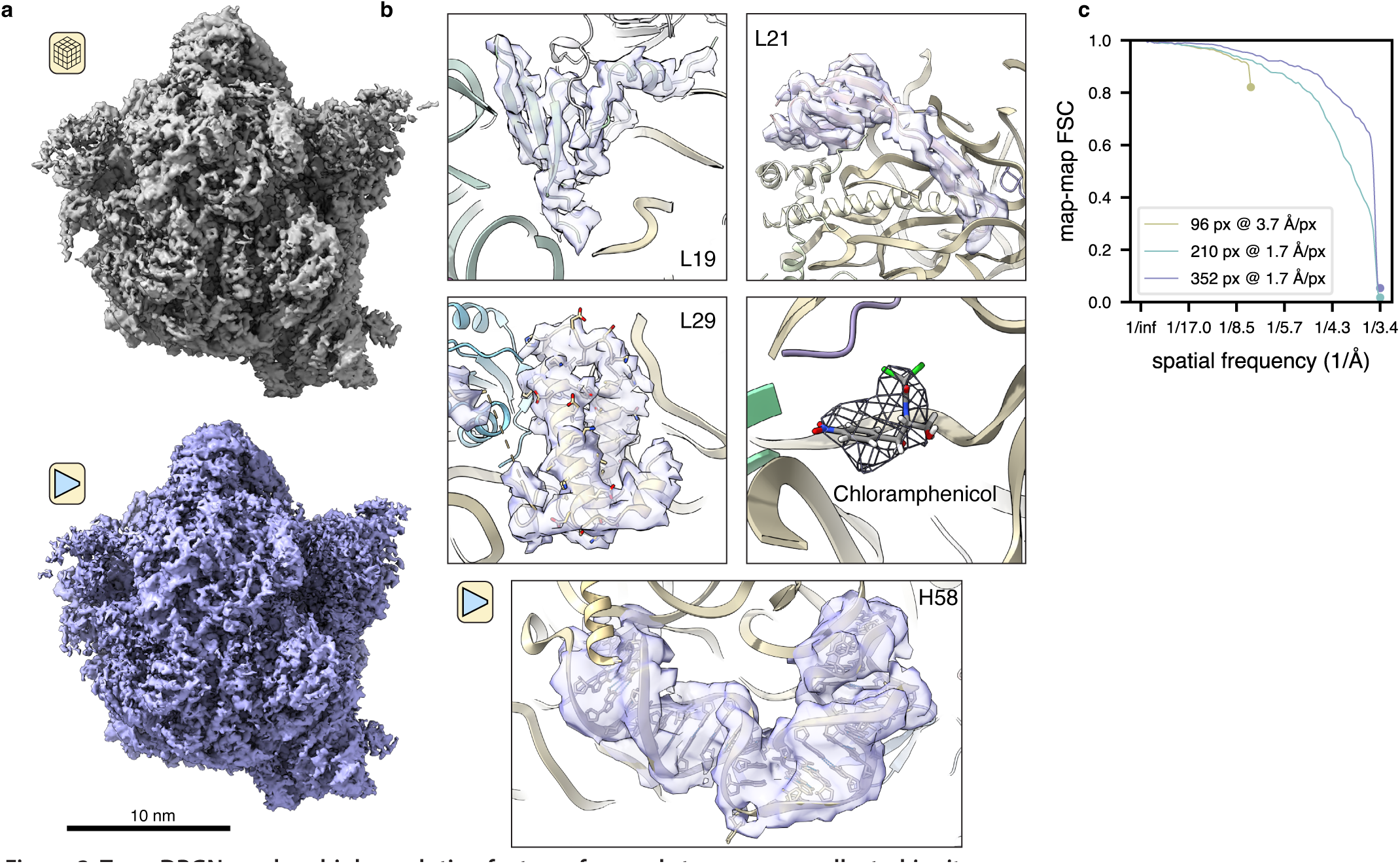
TomoDRGN resolves high resolution features from sub-tomograms collected *in situ*. **(a)** *M. pneumoniae in situ* ribosomal volume obtained from traditional STA processing (n=22,291 particles) (top) and tomoDRGN homogeneous reconstruction of the same particles (bottom). **(b)** Density maps from the tomoDRGN homogeneous reconstruction around indicated ribosome components. **(c)** Map-to-map Fourier Shell Correlations (FSC) of three tomoDRGN reconstructions of the particle stack in (a) extracted at indicated box and pixel sizes against corresponding STA volumes. Circles denote the Nyquist limit for each particle stack.

Encouraged by this result, we trained a heterogeneous tomoDRGN model on a down-sampled version of the particle stack and observed several distinct clusters in the resulting latent space (**Fig. 4a**, **left**). Generating volumes from these populated regions of latent space revealed that the majority of latent encodings corresponded to bonafide 70S ribosomes, as expected, whereas one subset corresponded to 50S ribosomal subunits, and another subset corresponded to apparent non-ribosomal particles (**Fig. 4a**, **right**). The non-ribosomal particles were further characterized by localizing them within each tomogram and providing them to RELION for *ab initio* reconstruction. Doing so suggested that most of these particles were false positive particle picks (**Extended Data Fig. 5**), highlighting tomoDRGN’s efficacy in sorting particles by structural heterogeneity generally, and in identifying errant particle picks specifically. We explored other approaches to separate 70S, 50S, and non-ribosomal particles, including using the trained tomoDRGN model to generate unique volumes corresponding to every particle’s latent embedding and either computing each volume’s similarity to the 70S STA map (**Fig. 4b**) or performing principal component analysis (PCA) in voxel space (**Fig. 4c**). Although these approaches produced results consistent with the clusters identified in latent space, for this dataset, the latent space clustering most clearly separated the 70S, 50S, and non-ribosomal particles.

**Figure 4:**
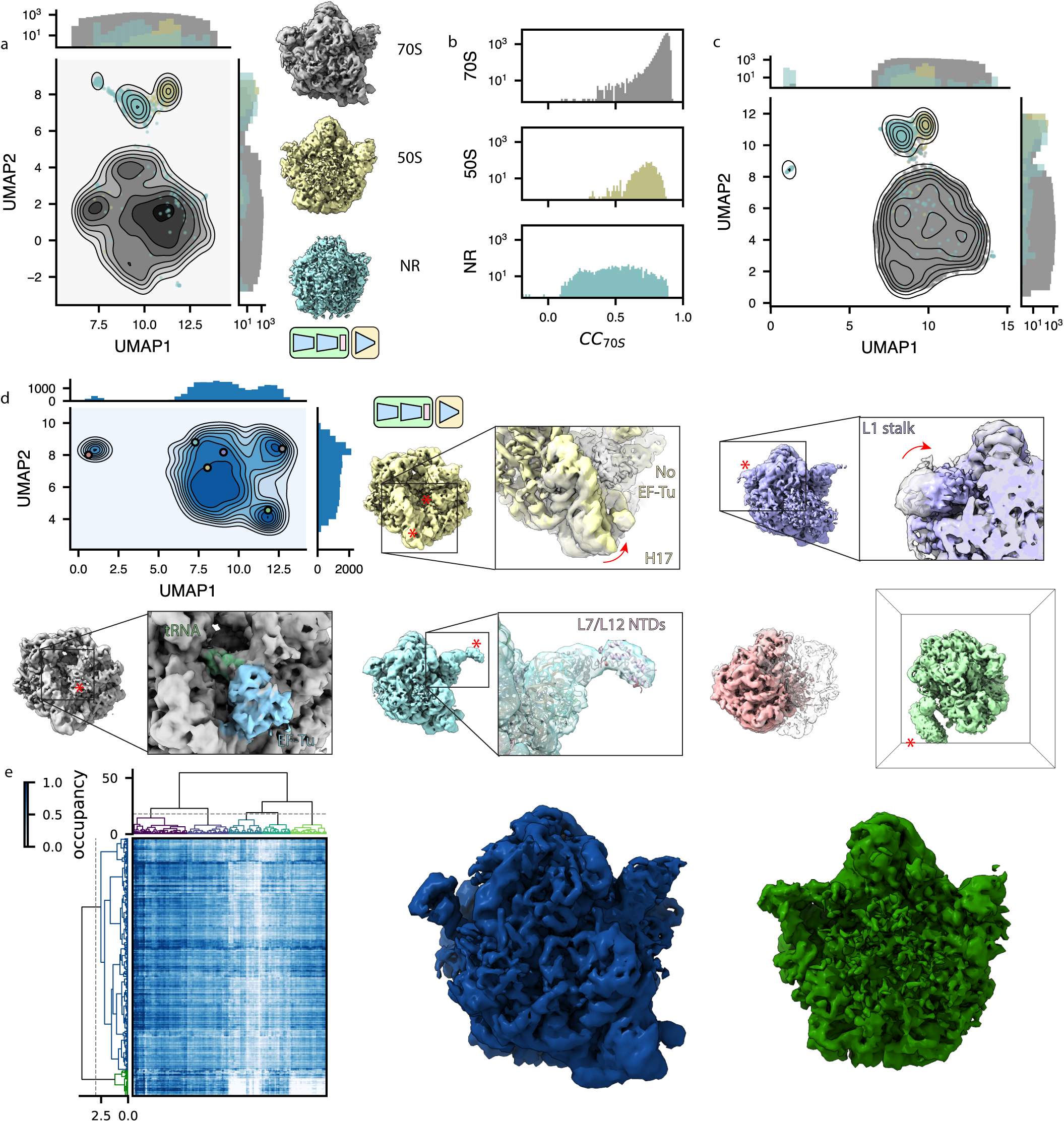
TomoDRGN uncovers structural heterogeneity in ribosomes imaged in situ. **(a)** UMAP of tomoDRGN latent embeddings (n=22,291 particles) shown as gray kernel density estimate (KDE), overlaid with scatter plot depicting latent embedding locations of large-ribosomal-subunit-only (yellow) or non-ribosomal particles (blue) identified via *k*=100 *k*-means classification of latent space and manual inspection of the 100 related volumes. Representative volumes generated from latent embeddings annotated as 70S, 50S, or non-ribosomal (NR) also depicted. **(b)** Volumes (box=96 px) were generated from every particle’s latent embedding, and volumetric cross-correlation (CC) between the 70S STA map and these volumes was calculated. Histograms of CC are shown for volumes assigned as 70S (top), 50S (middle) and non-ribosomal (bottom) particles as in (a). **(c)** Volumes from panel (b) were subjected to principal component analysis. UMAP dimensionality reduction of the first 128 principal components is plotted as a KDE with scatterplot corresponding to assignments of 70S, 50S, or non-ribosomal from (a) superimposed. **(d)** UMAP of tomoDRGN latent embeddings (n=20,981; non-ribosomal particles excluded). Colored volumes sampled from correspondingly colored points on UMAP plot are shown with red asterisks and insets highlighting regions of notable structural variability. A transparent grey volume corresponding to a tomoDRGN reconstruction of a 70S•EF-Tu volume is provided for visual reference. *(continued)* **(e)** MAVEn analysis (Sun *et al*., 2022) of 500 volumes sampled from the tomoDRGN model in panel (d) plotted as a clustered heatmap with columns corresponding to proteins and rRNA structural elements (Ward-linkage, Euclidean-distance), and rows corresponding to the 500 sampled volumes (Ward-linkage, Correlation-distance). Distinct volume classes corresponding to 50S and 70S particles as identified by a row-wise threshold on this clustermap are also shown.

Guided by the latent embeddings, we next filtered out the non-ribosomal particles and used this ‘clean’ subset to train a new heterogeneous tomoDRGN model. The resulting latent space and generated volumes revealed an array of structurally heterogeneous ribosomes (**Fig. 4d**). Prior analyses of this dataset focusing on translation cycle heterogeneity (Xue *et al*., 2022) identified a major class with the A- and P-tRNA binding sites occupied by tRNAs and several minor classes featuring variable occupancy and positions of tRNAs in the A and P sites and EF-Tu in the A site. Consistently, we observed that these states are highly represented in our sampled volumes, and we further observed additional conformational and compositional heterogeneity throughout the ribosome (**Supplemental Movie 1**). For example, we found a set of volumes lacking EF-Tu and with helix 17 of the 16S bent towards the now-unoccupied EF-Tu binding site. In other volumes, we observed pronounced motions of the L1 stalk. We also observed volumes with clear density for r-proteins L7/L12 in the expected 1:4 ratio of L10^CTD^:L7^NTD^/L12^NTD^ dimer of dimers. This observation was notable as this structural element is often unresolved in cryo-EM maps (Fromm *et al*., 2023; Stojkovic *et al*., 2020), likely due to this stalk’s dynamic nature and L7/L12’s ability to exchange off of the particle during purification (Chen *et al*., 2012). Observing this structure highlighted tomoDRGN’s ability to identify low abundance classes and emphasized the promise of the purification-free *in situ* structural analyses afforded by cryo-ET.

We next applied MAVEn (Kinman *et al*., 2023; Sun *et al*., 2022), which has previously been used to systematically interrogate the structural heterogeneity of volume ensembles guided by atomic models. Here, we observed a broadly uniform distribution of occupancies for all queried structural elements (*i.e.,* rRNA helices and r-proteins), with a notable exception of the 50S particle block, which lacks occupancy for any small subunit structural elements (**Fig. 4e**). We thus concluded that compositionally heterogeneous assembly intermediates were rare in this sample.

### TomoDRGN learns intermolecular heterogeneity

A grand promise of *in situ* cryo-ET is its potential to structurally characterize interactions between individual macromolecular complexes and their local environment (Tegunov *et al*., 2021; Turk and Baumeister, 2020). We hypothesized that tomoDRGN might perform well in this regard as its variational autoencoder architecture has a significant capacity to learn heterogeneity from the provided images, independent of the images being tightly or loosely cropped to the complexes under consideration. Indeed, our initial analysis revealed volume classes containing apparent intermolecular density truncated by the extracted box borders (**Fig. 4d**). To test tomoDRGN’s ability to analyze inter-complex structural heterogeneity, we extracted each ribosomal particle with a larger real-space box, effectively surveying the molecular neighborhood of each ribosome in the imaged cell. Training a new tomoDRGN model with these images revealed a similarly featured latent space with correspondingly diverse volumes (**Fig. 5a**). Many of the structures appeared to be disomes and trisomes, as previously reported (Tegunov *et al*., 2021), with measures of interparticle distance and the angular distribution to each ribosome’s nearest neighbor consistent with this interpretation (**Fig. 5b**). Intriguingly, the 50S population had an exceptionally broad distribution of nearest neighbor distances, and a subset of tomograms consisted almost exclusively of 50S ribosomes, whereas all other tomograms bore a more balanced distribution of all structural classes (**Fig. 5b-c**).

**Figure 5:**
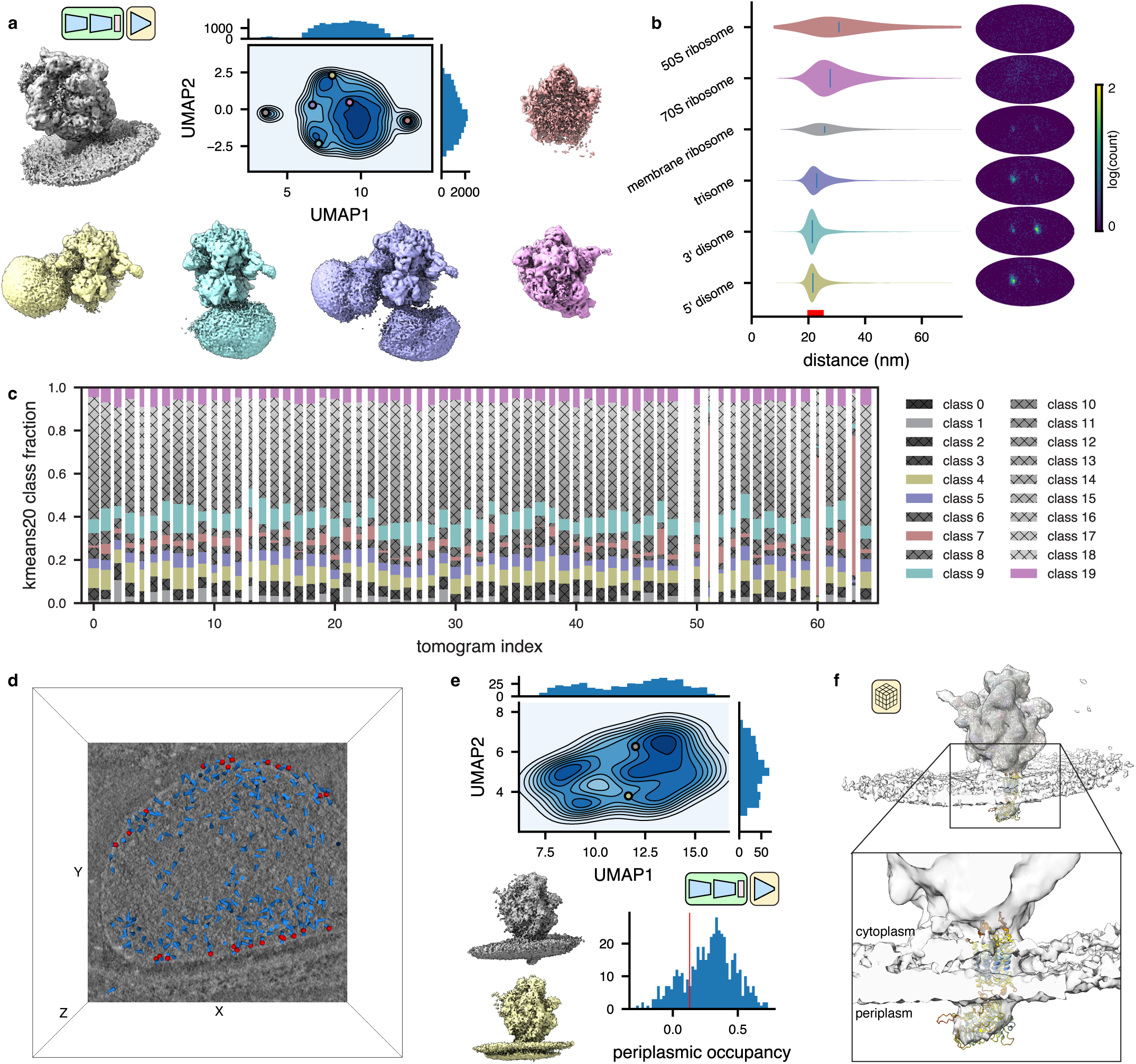
TomoDRGN captures intermolecular heterogeneity *in situ*. **(a)** UMAP of tomoDRGN latent embeddings of particles (n=20,981) re-extracted with box size ∼3x particle radius. Colored volumes sampled from correspondingly colored points in UMAP are shown. **(b)** Violin plot of the distance from each particle in the indicated classes from panel (a) to its nearest neighbor ribosome. Distribution colors are paired with those in (a). The right bound of the x-axis corresponds to the box diameter, and the red interval on the x-axis corresponds to typical inter-ribosome distances in a prokaryotic polysome. Mollweide projection histograms for each class highlighted in panel (a), depicts directions to each ribosome’s nearest neighbor ribosome, following rotation to the consensus pose. **(c)** Distribution of *k*=20 *k*-means classification of latent embeddings per tomogram. Column width is proportional to each tomogram’s fraction of the total particle count. Within a column, the height of each color is proportional to the population of that *k*-means class within that tomogram. Classes are colored as in (a). **(d)** Screenshot from tomoDRGN’s interactive tomogram viewer showing all ribosomes for a single tomogram (blue cones) with ribosomes corresponding to membrane-associated classes further annotated as red spheres. **(e)** UMAP of tomoDRGN latent embeddings (n=482) of membrane-associated ribosomes. Colored volumes are sampled from correspondingly colored points in latent space. Relative occupancy of globular periplasmic density (n=482) is plotted as a histogram with a red line noting manually assigned threshold defining particles bearing the periplasmic density (n=380). **(f)** STA reconstruction of membrane-associated ribosomes bearing periplasmic density identified by tomoDRGN with docked atomic model of *Mycoplasma pneumoniae* SecDF predicted using Alphafold (AF: A0A0H3DPH3).

Through this analysis, we observed a previously unreported ribosome structure with additional density corresponding to a lipid bilayer (**Fig. 5a**). To validate that this observed membrane density was not an artifact of the neural network, we mapped this set of apparently membrane-associated ribosomes to their original tomograms and observed that they exclusively corresponded to particles at the cell’s surface (**Fig. 5d**). To further identify residual heterogeneity within this group, we trained a new tomoDRGN model on this particle subset. We observed a relatively unfeatured latent space, with the majority (∼80%, as quantified by MAVEn), of sampled volumes bearing a flexible periplasmic density protruding from the membrane (**Fig. 5e**). Notably, we observed significant relative motion between the ribosome and the adjacent membrane, indicating that the ribosome is not held in rigid alignment with the membrane and holotranslocon during translocation (**Supplemental Movie 2**). Traditional STA on this periplasmic-positive subpopulation of ribosomes further resolved the periplasmic density, as well as smaller arches of density connecting the ribosome to the membrane (**Fig. 5f**, **Extended Data Fig. 6c**). Rigid body docking using atomic models of likely transmembrane protein complexes into this density supported that we had identified ribosomes bound to SecDF, a subcomplex of the Sec holotranslocon with a relatively large extracellular globular domain encoded in the *M. pneumoniae* genome (**Fig. 5f**). This result highlighted the efficacy of tomoDRGN’s iterative particle curation and refinement approach in unveiling new structures buried in highly heterogeneous *in situ* datasets.

## DISCUSSION

In this work, we introduce tomoDRGN, which, to our knowledge, is the first neural network framework capable of simultaneouslymodelingcompositionalandconformational heterogeneity from cryo-ET data on a per-particle basis. TomoDRGN achieves this using a bespoke deep learning architecture and numerous accelerations designed to exploit redundancies inherent to cryo-ET data collection. We note that the major heterogeneity analyses demonstrated in this manuscript were also tested with cryoDRGN (Zhong *et al*., 2021). However, cryoDRGN ultimately did not match tomoDRGN’s performance on cryo-ET data as it incorrectly classified simulated data, predominantly learned non-biological structural heterogeneity, and produced highly variable latent embeddings and volumes for different tilt images of the same particle (**Extended Data Figs. 7-9**). We note that an alternative approach of mapping single sub-tomogram volumes to single latent coordinates would theoretically function within the cryoDRGN framework but would: 1) be less computationally tractable due to cubic scaling of the number of voxel coordinates to be evaluated per particle; and 2) may be predisposed towards learning heterogeneity driven by missing wedge artifacts common to sub-tomogram volumes.

Other tools to explore conformational heterogeneity from a cryo-ET dataset exist (Bharat and Scheres, 2016; Harastani *et al*., 2021; 2022; Himes and Zhang, 2018). However, they each rely on some degree of imposed prior knowledge, either in the form of “mass conservation” to describe continuous changes from a consensus structure, which is often derived from a provided atomic model (Harastani *et al*., 2021; 2022); assumptions of linear relationships between structures (Himes and Zhang, 2018); or the assertion that a small number of discrete structures exist (Bharat and Scheres, 2016). In contrast, the cryoDRGN/tomoDRGN approaches provide a greater degree of generality that we have found enables largely unsupervised learning of highly complex combinations of compositional and continuous conformational heterogeneity. Given the extent of structural heterogeneity observed with cryoDRGN in single particle datasets using purified samples (Sekne *et al*., 2022; Vasyliuk *et al*., 2022), we expect tomoDRGN to uncover similar structural variation within a rapidly expanding set of samples imaged *in situ* with cryo-ET.

In addition to characterizing the structural heterogeneity of isolated particles, we expect that tomoDRGN’s ability to reanalyze particle stacks at different spatial scales (*i.e.*, different real space box sizes) will prove widely useful in correlating intramolecular structural changes with structural variability in areas adjacent to the particle (**Fig. 6**). For example, when analyzing isolated ribosomal particles in EMPIAR-10499, we identified a set of particles decoding EF-Tu•tRNA and, when analyzing particles in a larger spatial context, we found a set of ribosomes associated with the cell membrane. A straightforward comparison enabled calculating the co-occurrence of these properties, revealing that approximately 13% of the membrane-associated ribosomes also appear to be decoding EF-Tu•tRNA. Of particular note, tomoDRGN allows us to generate a unique 3-D volume corresponding to each particle’s latent embedding. We anticipate this approach will allow researchers to populate low SNR tomograms with particle-specific density maps at approximately nanometer resolution and to explore the resultant spatial distributions of heterogeneous structures.

**Figure 6:**
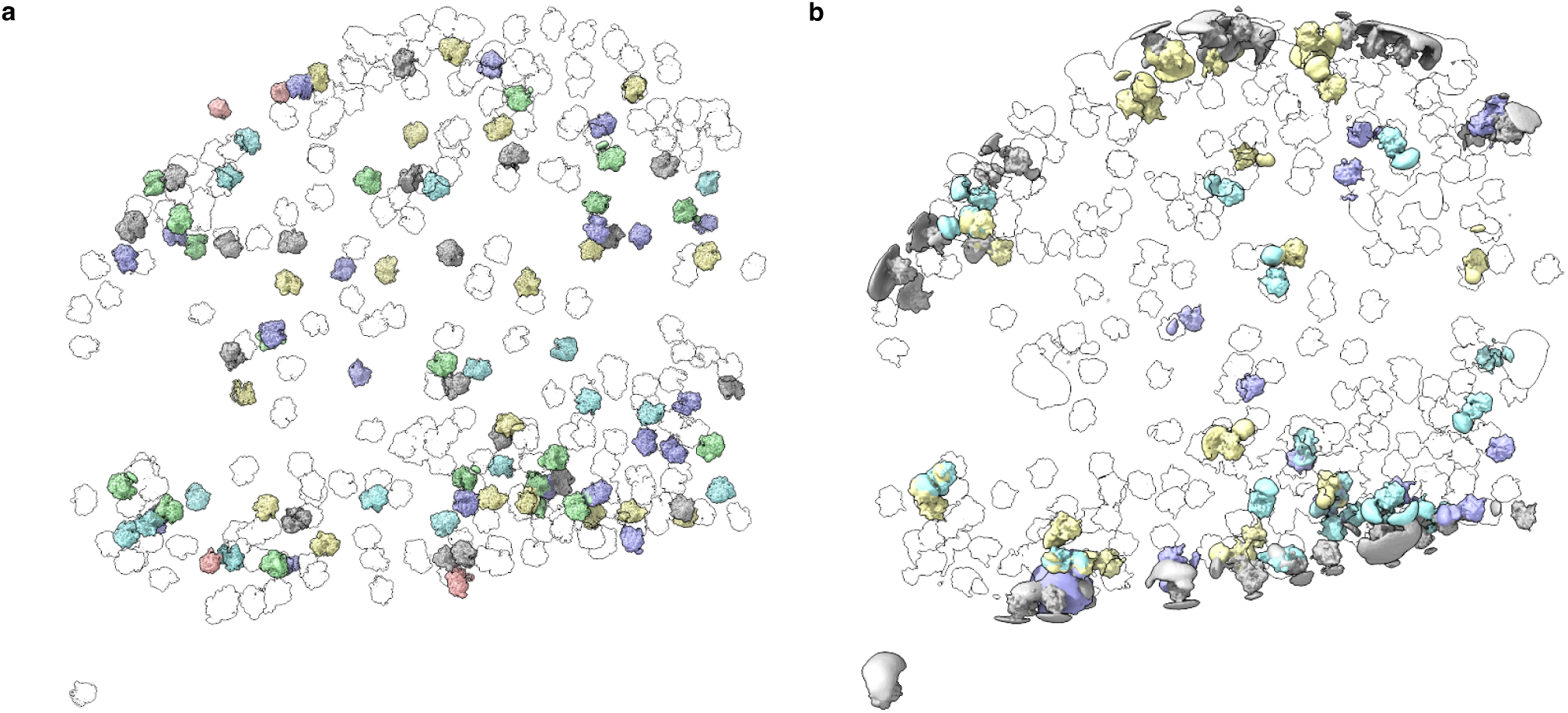
TomoDRGN visualizes structurally heterogeneous macromolecular complexes with spatial context. **(a)** Ribosomes from one EMPIAR-10499 tomogram rendered in ChimeraX. Volumes were generated for each ribosome using tomoDRGN and colored according to the intramolecular latent classification shown in Fig. 4d, and positioned correspondingly within the reconstructed cell. Transparent ribosomes correspond to *k*=20 *k*-means classes not highlighted in Fig. 4d. **(b)** Ribosomes from the same tomogram depicted in (a) rendered in ChimeraX. Volumes were generated for each ribosome and colored according to the intermolecular latent classification shown in Fig. 5a, and positioned correspondingly within the reconstructed cell. Transparent ribosomes correspond to *k*=20 *k*-means classes not highlighted in Fig. 5a.

Finally, the analyses enabled by tomoDRGN are inherently iterable. Our initial tomoDRGN analysis of EMPIAR-10499 revealed a population of non-ribosomal particles that we had failed to filter with traditional classification-based approaches. Excluding such particles and retraining at multiple spatial scales resolved intra- and inter-molecular structural heterogeneity, and retraining exclusively on a subset of membrane-associated ribosomes identified extracellular density that likely corresponded to the SecDF subcomplex. Given that tomoDRGN has the potential to identify many such distinct classes, we encourage users to embrace this iterative, branching approach. Some recently introduced software packages (Rice *et al*., 2022; Tegunov *et al*., 2021) explicitly support such “molecular sociology” where co-refinement of multiple distinct structures derived from a common data source enables global improvement of map quality. We anticipate tomoDRGN will form a virtuous cycle when interfacing with such software.

## MATERIALS AND METHODS

### TomoDRGN design and software implementation

#### General architecture

TomoDRGN is forked from cryoDRGN, and although we summarize the core aspects of the method here, readers are pointed to related cryoDRGN publications for further details (Bepler *et al*., 2019; Kinman *et al*., 2023; Zhong *et al*., 2021; Zhong *et al*., 2019). In brief, tomoDRGN is a variational autoencoder (VAE) with encoder and decoder networks comprised of multi-layer perceptrons (MLPs). TomoDRGN’s encoder learns a function (*E*) to map a set of *j* distinct tilt images (size × pixels) of particle *i* to a low dimensional latent encoding *z_i_* of dimension *z*; that is, : . The encoder MLP is comprised of two sub-networks that process *j* tilt images for each particle as follows. First, the 2-D Hartley transform of each tilt image is passed separately through encoder A to produce a set of *j* intermediate encodings. These *j* intermediate encodings are then pooled and passed together through encoder B to output the particle’s final latent embedding *z_i_*. The pooling step concatenates intermediate encodings along the tilt image axis by default, but also supports operations such as *max* and *mean*, which are inherently permutation-invariant. All experiments presented here concatenate the intermediate encodings.

TomoDRGN’s decoder follows from that of cryoDRGN (Zhong *et al*., 2021), and uses a Gaussian featurization scheme for positional encoding in Fourier space (Tancik *et al*., 2020) as follows. Spatial coordinates are normalized to span [-0.5, 0.5] in each dimension, and a (fixed) positional encoder transforms each spatial coordinate to a basis set of D sinusoids with frequencies sampled from a scaled standard latent variable, the decoder predicts the corresponding voxel intensity as ρ_θ_(*V|k, z*). Applying the Fourier Slice Theorem (Bracewell, 1956), 3-D Fourier coordinates corresponding to 2-D projection image *X_i_* are derived by rotating a 2-D lattice by the orientation of the volume *V_i_* during imaging. Given a fixed latent coordinate sampled from *q*ξ(*z_i_*|*X_i_*) and the posed coordinate lattice, the reciprocal space image is reconstructed pixel-by-pixel via the decoder ρ_θ_(*V|k, z_i_*). The reconstructed image is then translated in-plane and multiplied by the CTF. The negative log-likelihood of the image is then computed as the mean squared error between the input and reconstructed image. The optimization function is the sum of the image reconstruction error and the KL divergence (KLD) of the latent encoding:

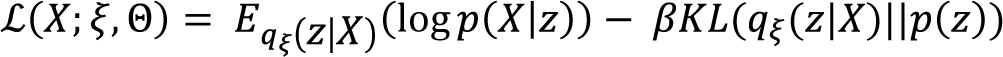

In this equation, the regularizing KLD term is weighted by β, which is set to 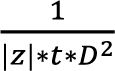 where D is the box size, t is the number of tilts, and |*z*| is the dimensionality of the latent space.

#### Lattice masking and reconstruction weighting

Critical dose is calculated for each spatial frequency using an empirical exposure-dependent amplitude attenuation curve derived for cryo-EM data (Grant and Grigorieff, 2015). The optimal dose is approximated to 2.51284 × as in the original study (Grant and Grigorieff, 2015; Hayward and Glaeser, 1979). Spatial frequencies (coordinates) of a tilt image exceeding the corresponding optimal doses are excluded from decoder network evaluation and loss calculation by a lattice mask during network training. Following error calculation of the input image against axis, where D is the box size of an input image, and *feat_ sigma* is set to 0.5. These positionally encoded coordinates, concatenated with the *z*-D latent coordinate, are then passed to the decoder; that is, in totality, *D*:ℝ^3+*z*^ → ℝ Unless otherwise specified, models were trained for 50 epochs with batch size 1 (particle), AdamW optimizer with learning rate 0.0002, and weight decay 0.

#### Training system

Input images are modeled as linear 2-D projections of 3-D volumes, convolved by the contrast transfer function (CTF), with externally-derived rotation, translation, and CTF parameters. Heterogeneity among volumes is modeled via a continuous latent space sampled by a latent variable z per particle. The latent encoding for a given image is taken as the maximum *a posteriori* of a Gaussian distribution parameterized by outputs from the encoder network, μ_*z*|*X*_ and Σ_*Z|X*_, whereas the prior on the latent distribution is a standard normal distribution *N*(0, ***I***). Thus, the variational encoder *q*ξ(*z*|*X*) produces a variational approximation of the true posterior *p*(*z|X*).

The coordinate-based decoder models structures in reciprocal space: given a spatial frequency *k* ϵ ℝ^3^ and a the reconstructed and CTF-weighted voxels, the squared differences are weighted (1) per-frequency by the exposure dependent amplitude attenuation curve (a function of tilt image index and spatial frequency), and (2) globally by the cosine of the stage tilt angle (a function of tilt image index). This weighted reconstruction error is backpropagated accordingly.

### Random tilt sampling

During dataset initialization, the number of tilt images per particle is parsed via the rlnGroupName star file column using the syntax in Warp/M of *tomogramID_particleID*. The minimal number of tilt images present for any particle is then stored as the number of images to be sampled from each particle during network training and evaluation (this value also sets the input dimensionality of encoder B when using concatenation pooling). By default, sampling is performed randomly without replacement per-particle, and the subset and ordering of sampled tilts is updated each time a particle is retrieved during training or evaluation.

### Simulated dataset generation

Cryo-ET data simulation was performed using scripts in the cryoSRPNT (<u>cryo</u>-EM <u>S</u>imulation of <u>R</u>ealistic <u>P</u>articles via <u>N</u>oise <u>T</u>erms) GitHub repository. Density maps of four assembly states of the bacterial 50S ribosome (classes B - E) were obtained from EMD-8440, EMD-8441, EMD-8445, and EMD-8450, respectively (Davis *et al*., 2016). The *project3d. py* script was used to create noiseless projections of each volume as follows. First 5,000 random particle poses were sampled over SO(3). Each randomly posed particle was then rotated following a dose-symmetric tilt series scheme from 0° to ±60° with 3° steps in groups of 2 over 41 tilts, resulting in a total of 205,000 unique poses per volume. Each posed volume was projected along the z-axis to create noiseless images.

The *acn.py* script was used to corrupt the noiseless projections using a standard cryo-EM image formation model (Baxter *et al*., 2009) augmented by tilt-series specific subroutines as follows. First, noiseless projections were Fourier-transformed, dose-weighted following an empirical exposure dependent amplitude attenuation curve at 3 e^-^/Å^2^/tilt to simulate SNR decrease due to radiation damage (Grant and Grigorieff, 2015), and inverse Fourier-transformed. Structural noise was added with an SNR of 1.4, and particles were then weighted by cosine(tilt) to simulate SNR decrease due to increased sample thickness. Projections were then convolved with the 2-D CTF with defocus values sampled from a mixture of Gaussian-distributed defoci with means between -1.5 µm to -3.5 µm in 0.5 µm steps and a standard deviation of 0.3 µm. Other CTF parameters included no astigmatism, 300 kV accelerating voltage, 2.7 mm spherical aberration, 0.1 amplitude contrast ratio, and 0° phase shift. Finally, shot noise was added with a SNR of 0.1, for a final SNR of 0.05. Particle stacks of each class were Fourier cropped to box sizes of 256px (bin1), 128px (bin2), and 64px (bin4).

### TomoDRGN network training on simulated data

TomoDRGN homogeneous network training was performed on the 5,000 simulated class E particles. TomoDRGN heterogeneous network training was performed on all 20,000 simulated particles from classes B-E. Unless otherwise specified, figures illustrate results on the bin2 datasets, with network architectures summarized as *nodes_ per_layer* x *layers* as follows: of 128×3 (encoder A), 128×3 (encoder B), and 256×3 (decoder). The dimensionality of the intermediate encoding was 32 and that of the final latent encoding was 128. Each model was trained utilizing dose and tilt loss weighting, dose frequency masking, and random tilt sampling, unless specified otherwise. Classification was performed directly on the latent embeddings with *k*=4 *k*-means clustering as implemented in scikit-learn. The dataset’s latent value nearest each *k*-means cluster center was used to generate a 3-D volume representative of that cluster.

### Sub-tomogram averaging of EMPIAR-10499 ribosomes

Raw tilt movie data was downloaded from EMPIAR-10499. Movies were aligned and initial CTF estimation was performed in Warp (Tegunov and Cramer, 2019) as previously (Tegunov *et al*., 2021). Automated fiducial-based tilt series alignment was performed using dautoalign4warp (Burt *et al*., 2021) within the Dynamo package running in a Matlab environment (Castano-Diez *et al*., 2012). Alignment parameters were then used to generate tomograms at 10 Å/px in Warp. Template matching was performed in Warp using a 40 Å lowpass filtered ribosome volume generated from manually picked particles, keeping particles with a minimum separation of 80 Å (974,804 particles). The top 3% of particles by figure-of-merit across all tomograms were kept (29,245 particles). Sub-tomograms were extracted in Warp at 10 Å/px. *Ab initio* model generation and 3-D refinement were performed in RELION 3.1 (Bharat and Scheres, 2016) resulting in a density map with Nyquist-limited resolution. Sub-tomograms were re-extracted in Warp at 4 Å/px for further RELION 3-D refinement and 3-D classification with *k*=4 classes to remove false positive particle picks. The remaining 22,291 ribosomal particles were refined to a nominal resolution of 8.1 Å. Between each round of refinement and classification, particles were deduplicated in RELION with a cutoff distance of 80Å (removing a total of 360 particles throughout processing). The final 22,291 particles were imported to M and to produce a 3.5 Å resolution map as reported previously (Tegunov *et al*., 2021). Particles were then exported as image series sub-tomograms from M at several pixel and box sizes for tomoDRGN training, including three “single ribosome diameter” scales: 96 px at 3.71 Å/px, 210 px at 1.71 Å/px, 352 px at 1.71 Å; and one “multiple ribosome diameter” scale: 200 px at 3.71 Å/px. Particles were also exported as volume series sub-tomograms using M at 64 px 6 Å/px and 192 px 4 Å/px for validation of tomoDRGN heterogeneity analysis with traditional STA tools (see below) and for generation of requisite metadata for mapping particles to tomogram-contextualized locations in the tomoDRGN analysis Jupyter notebook.

### TomoDRGN network training on EMPIAR-10499

TomoDRGN homogeneous network training was performed on the 22,291 image series particles extracted at each of the “single ribosome diameter” image series sub-tomograms described above, or on select subsets at 96 px at 3.71 Å/px for interrogating heterogeneity in specific particle subsets. Unless specified otherwise, the network architecture was 512×3 (decoder). Each model was trained utilizing dose and tilt loss weighting, dose frequency masking, and random tilt sampling.

TomoDRGN heterogeneous network training was performed on the same stack of 22,291 image series particles at box 96 px and 3.71 Å/px. Unless specified otherwise, the network architecture was 256×3 (encoder A), 256×3 (encoder B), and 256×3 (decoder) with the dimensionality of the intermediate encoding set to 128, and that of the final latent encoding set to 128. Each model was trained utilizing dose and tilt loss weighting, dose frequency masking, and random tilt sampling. Classification was performed directly on the latent embeddings with either *k*=20 (used for general visualization) or *k*=100 (used for detailed visualization and particle filtering) *k*-means clustering as above. The dataset’s latent value nearest each *k*-means cluster center was used to generate a 3-D volume representative of that cluster. Following exclusion of 1,310 non-ribosomal particles by separation of such volumes from *k*-100 classification, the remaining 20,981 particles were used to train new tomoDRGN models at box sizes of 96 and 200 px with 3.71 Å/px sampling. Membrane associated ribosomes (482) identified by *k*-100 classification of the 200 px trained dataset were further isolated to train a new tomoDRGN model with the parameters noted as above.

### Visualization and validation

#### Python scripts

A number of Python scripts were generated to quantify various properties of tomoDRGN outputs. Classification accuracy of tomoDRGN latent encodings learned for simulated datasets was evaluated by generating a confusion matrix (**Fig. 2e**). Classification reproducibility was evaluated for 100 randomly initialized classifications by calculating the Adjusted Rand Index (ARI) (Hubert and Arabie, 1985) (**Extended Data Fig. 7f**). The ARI measures a label-permutation-invariant similarity between two sets of clusterings and scales from 0 (random labeling) to 1 (identical labeling). Here, we used ARI to measure the similarity between the tomoDRGN or cryoDRGN latent clusters and the ground truth class labels.

Volume consistency was assessed by calculating the map-map correlation coefficient (Real space) and map-map FSC scripts available within the tomoDRGN software. Before calculating map-map FSC curves, a soft mask was calculated and applied in Real space. Masks were defined by binarizing the map at ½ of the 99^th^ voxel intensity percentile, dilating the mask by 3 px, and softening the mask using a falling cosine edge applied over 10 px. For computational efficiency, in some instances, the map-map correlation coefficient metric (CC) (Afonine *et al*., 2018) was used to quantify map-to-map similarity.

Heterogeneity of a set of EMPIAR-10499 pre-filtered ribosome volumes generated by tomoDRGN was quantified by generating all volumes from the final epoch of training’s latent values and either (1) calculating the map-map CC to the STA 70S map for each tomoDRGN volume (**Fig. 4b**), or (2) performing principal component analysis on the array of all volume’s voxels (shape *n_volumes_* × *D*^3^) followed by UMAP dimensionality reduction of the first 128 principal components (**Fig. 4c**).

Finally, Python scripts were used to identify each particle’s nearest neighbor in each tomogram, calculate the distance to the nearest neighbor, and calculate the angle to the nearest neighbor after rotating to the STA consensus reference frame (**Fig. 5c**).

#### Volume subset validation

Subsets of the EMPIAR-10499 ribosomes were identified by tomoDRGN as non-ribosomal (n=1,310), 50S (n=852), 70S (n=20,129), or membrane-associated (n=482). Non-ribosomal particles were reprocessed in RELION 3.1 using *ab initio* volume generation with *k*=5 volume classes and all other parameters at their defaults. The 50S, 70S, and membrane-associated ribosome populations were reprocessed in RELION 3.1 using 3-D refinement against a corresponding real-space cropped 70S volume lowpass filtered to 60 Å. The same three particle subsets were also used to train tomoDRGN homogeneous networks as an additional validation, with identical training parameters to the full particle stack training detailed above.

#### Visualization of tomoDRGN volumes in situ

The subtomo2chimerax script (https://zenodo.org/ <u>record/6820119</u>) was adapted to handle tomoDRGN’s unique sub-tomogram volumes per particle and is implemented in tomoDRGN. This script places each particle’s volume at its source location and orientation in the tomogram context using ChimeraX for visualization (Goddard *et al*., 2018; Pettersen *et al*., 2021). All volumes corresponding to EMPIAR-10499 tomogram 00256 were generated by tomoDRGN at box size 64 px and 5.55 Å/px using latent coordinates from tomoDRGN models in **Fig. 4d** and **Fig. 5a**, and placed in tomogram 00256 with coordinate and angle values extracted from the STA refinement in M.

### Atomic model-guided analyses

To aid interpretation of tomoDRGN density maps, atomic models of the 70S ribosome (7PHA, 7PHB, and 4V89 which highlighted the L7/L12 dimers) were docked into density maps as rigid bodies using ChimeraX. The rRNA of 7PHB was segmented into distinct chains corresponding to rRNA helices (Petrov *et al*., 2014) following the MAVEn protocol (Kinman *et al*., 2023) for model-based analysis of volume ensembles (https://github.com/lkinman/MAVEn). The predicted atomic model for *M.pneumoniae* SecDF was downloaded from AlphaFold (ID: A0A0H3DPH3) and docked into the membrane-associated ribosome STA map in ChimeraX as a rigid body. Other components of the canonical Sec holotranslocon and oligosaccharyltransferases were either absent in the *M. pneumoniae* genome or lacked the observed extracellular domain.

### CryoDRGN network training

CryoDRGN v0.3.4 was used to train models for both the simulated ribosome dataset (n=20,000) and the unfiltered EMPIAR-10499 dataset (n=22,291), using corresponding simulated or STA-derived poses and CTF parameters. Because cryoDRGN treats each input image independently, each dataset was reshaped to collapse the tilt axis dimension, resulting in particle stacks of size n=820,000 and n=913,931, respectively. Networks were trained with architecture 128×3 or 128×6 (encoder), latent dimensionality 8 or 128, and 256×3 (decoder), as annotated. All models were trained with hyperparameters intended to maximize similarity to the respective tomoDRGN analysis: batch size 40, gaussian positional featurization, 50 epochs of training, automatic mixed precision enabled, and all other parameters adopting default values. Latent space classification and volume sampling were performed as described for tomoDRGN above.

### Performance benchmarking

All tomoDRGN and cryoDRGN models were trained on a cluster with nodes each equipped with 2x Intel Xeon Gold 6242R CPU (3.10 GHz, 512 GB RAM) and 2x Nvidia GeForce RTX 3090. Reported training times may in some cases be overestimates as up to two jobs were allowed to train or evaluate simultaneously on the same node.

## SUPPLEMENTARY MOVIES

**Supplemental Movie 1: Structural heterogeneity in the large ribosomal subunit.**

Volumes were sampled from the tomoDRGN model in Fig. 4d using k=100 k-means clustering of latent space. Density for the 30S was removed using the Volume Zone feature in ChimeraX, guided by atomic model 7PHB, to reveal distinct conformation and compositional states of the large subunit. Note conformational and compositional heterogeneity in tRNA and elongation factor binding sites, which are found along the midline of the particle.

**Supplemental Movie 2: Membrane-associated ribosomes exhibit flexible attachment.**

Volumes were generated for all particles used to train the model in Fig. 5d. The tertile of volumes with highest SecDF occupancy are displayed, ordered by increasing occupancy (n = 162). Note significant dynamics in the orientation of the membrane relative to the associated ribosome.

## Supporting information

Supplemental Video 1: Structural heterogeneity in the large ribosomal subunit.

Supplemental Video 2: Membrane-associated ribosomes exhibit flexible attachment.

## ACKNOWLEDGEMENTS

We thank Laurel Kinman and Ellen Zhong for helpful discussion and feedback, and the MIT-IBM Satori team and the MIT SuperCloud Supercomputing Center for HPC computing resources and support. This work was supported by NIH grants R01-GM144542, 5T32-GM007287, and NSF-CAREER grant 2046778, the Sloan Foundation and the Whitehead Family.

## AUTHOR CONTRIBUTIONS

BMP and JHD conceived the work. BMP implemented the tomoDRGN method. BMP and JHD designed the experiments. BMP performed and analyzed the experiments. BMP and JHD wrote the manuscript.

## DATA AVAILABILITY

Extracted particle sub-tomograms from EMPIAR-10499 will be deposited to EMPIAR. The membrane-associated ribosome from EMPIAR-10499 generated by RELION will be deposited to EMDB. The trained tomoDRGN and cryoDRGN models used to analyze EMPIAR-10499 will be deposited at zenodo.org. Simulated data and corresponding trained tomoDRGN and cryoDRGN models are available upon request.

## SOFTWARE AVAILABILITY

TomoDRGN is distributed as free and open-source software under the GPL-3.0 license. Source code, installation instructions, and example usage are available at <u>https://</u> <u>github.com/bpowell122/tomodrgn</u>. Version 0.2.2 was used in this study. Scripts used to generate simulated data are available at https://github.com/bpowell122/cryoSRPNT. Version 0.1.0 was used in this study.

## EXTENDED DATA FIGURES

**Extended Data Figure 1:**
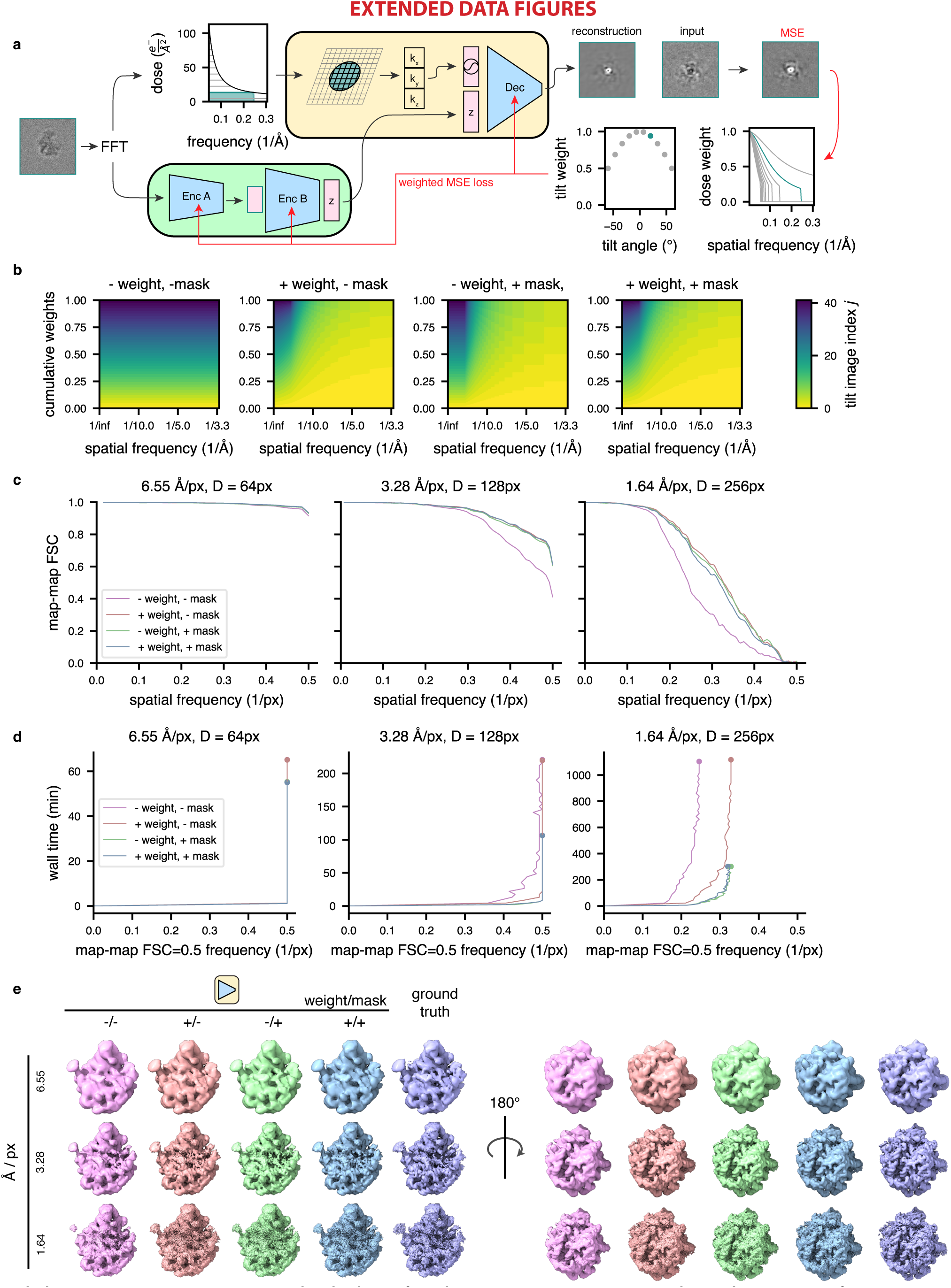
Training on a weighted subset of pixels improves reconstruction quality and compute performance. **(a)** Graphical overview of the dose filtering scheme (applied upstream of the decoder) and dose and tilt weighting scheme (applied during reconstruction error calculation) for a single representative tilt image. Filtering: the fixed optimal exposure curve is used to determine which spatial frequencies will be considered as a function of dose; the decoder processes only Fourier lattice coordinates within this mask (green lattice circle). Weighting: the squared error of the reconstructed Fourier slice is weighted per-frequency by the exposure-dependent amplitude attenuation curve and per-slice by the cosine of the corresponding stage tilt angle, before mean reduction and backpropagation (red arrows). **(b)** Relative weight of each tilt image assigned to a particle’s reconstruction error during model training as a function of spatial frequencies (x-axis), and tilt and dose, which are colored yellow to blue from low-to-high dose and tilt angle, assuming a dose symmetric tilt scheme (Hagen *et al*., 2017). **(c)** Map-map FSC of simulated class E large ribosomal subunit volumes (Davis *et al*., 2016) compared to tomoDRGN homogeneous network reconstructions in the presence or absence of the weighting or masking schemes at varying box and pixel sizes. **(d)** Spatial frequencies corresponding to FSC=0.5 map-map correlation with the ground truth volume plotted against wall time for model training. **(e)** Final tomoDRGN reconstructed volumes (left and center) and ground truth volumes (right) in the presence or absence of the weighting or masking schemes at box and pixel sizes assessed in panels (c) and (d).

**Extended Data Figure 2:**
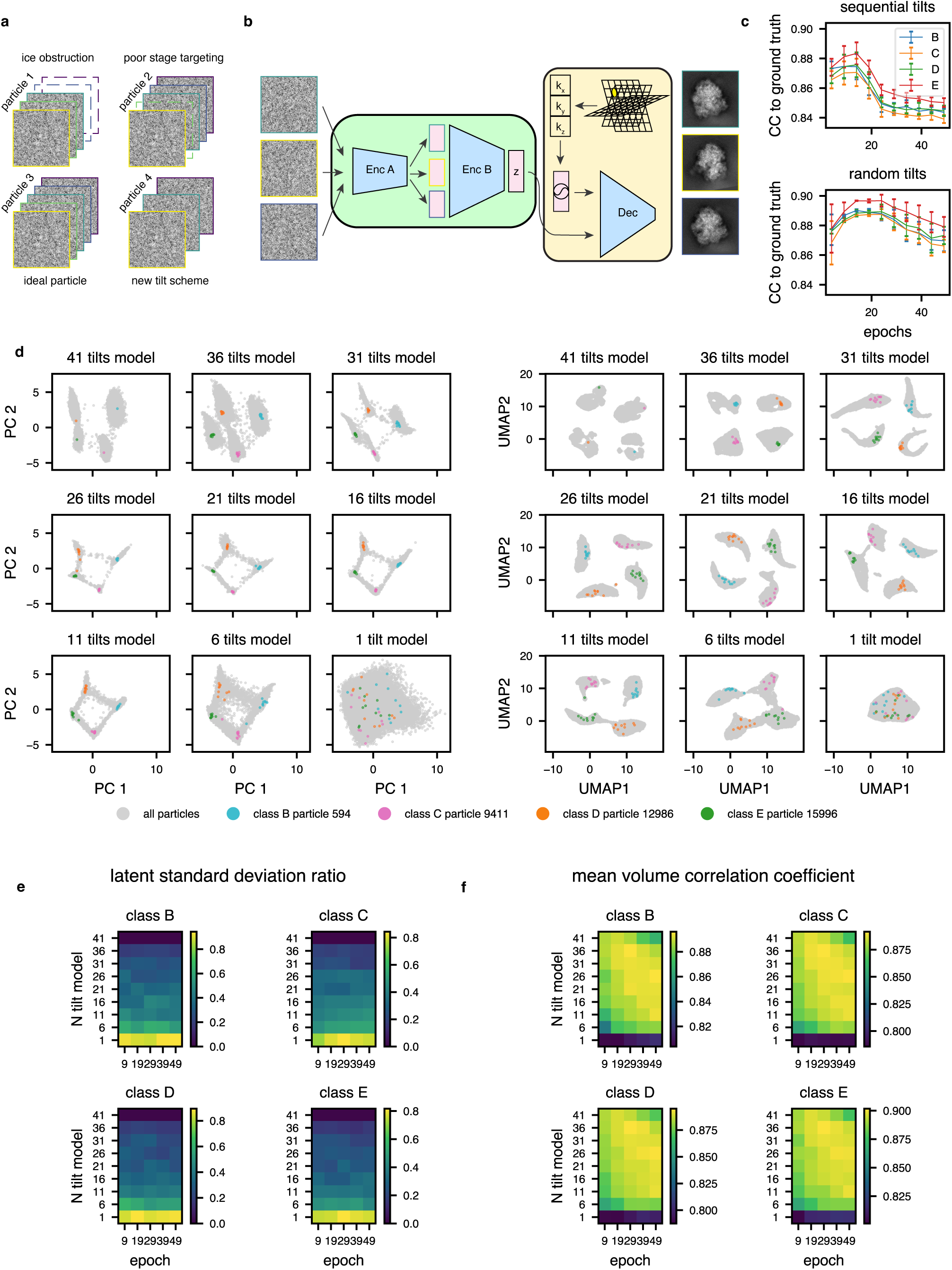
Random per epoch tilt selection enables flexible and robust model training for datasets with non-uniform numbers of tilt-images per particle. **(a)** Graphical summary of a dataset with non-uniform numbers of tilt images per particle. Here, the minimum number of tilt images for any particle is 3. **(b)** Corresponding tomoDRGN network architecture for random sampling and ordering of 3 tilt images per particle. **(c)** Mean per-class volumetric correlation coefficient for identical tomoDRGN models trained on 41 sequentially sampled tilts (top) or 41 randomly sampled tilts (bottom). At 5 epoch intervals, 25 random volumes were generated from each class for correlation coefficient calculation to ground truth ribosome assembly intermediate volumes (classes B-E). Error bars denote standard error of the mean CC. **(d)** Nine tomoDRGN models with identical architectures were trained with the indicated number of tilts sampled per particle (total available tilts = 41). PCA (left) and UMAP (right) dimensionality reduction of each final epoch’s latent embeddings. Once trained, up to 10 randomly sampled and permuted tilt images for one representative particle from each volume class were embedded using the corresponding pretrained tomoDRGN model and are superimposed as colored points. Note increased dispersion of colored points as number of tilts sampled during training decreased. **(e)** For each ribosomal large subunit class (B-E), 25 particles were randomly selected and up to 10 subsets of their tilt images were randomly sampled and permuted as in (d). In the heatmap, row indices refer to models trained in (d) using different numbers of sampled tilts (1-41), and columns denote epochs of training with that model. For each particle, each tilt subset was evaluated with the corresponding tomoDRGN model and the ratio of standard deviations of each particle’s 10 latent embeddings to all particles’ latent embeddings was calculated. The mean ratio across all particles, which measures the dispersion of encoder embeddings, is plotted per ribosomal LSU class. Here, lower dispersion indicates better performance. **(f)** Particles and tilt subsets were selected as in (e). At each indicated epoch of training, the corresponding tomoDRGN model was used to generate volumes for each particle’s tilt subsets. For each such volume, the correlation coefficient was calculated between that volume and the corresponding ground truth volume. The mean across all particles at each epoch for each model is shown as a heatmap per ribosomal LSU class. Here, higher CC indicates improved performance.

**Extended Data Figure 3:**
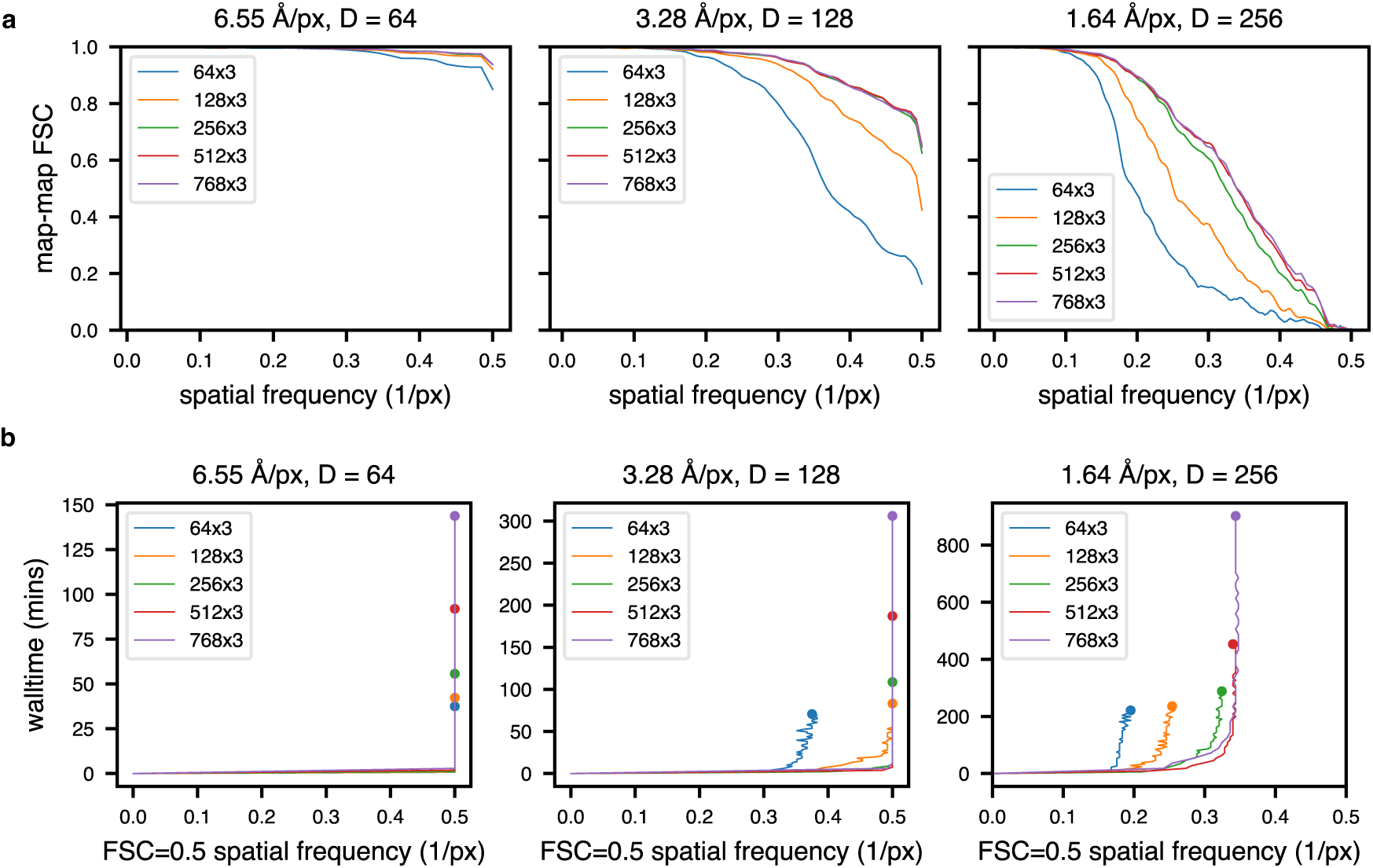
TomoDRGN training statistics for homogeneous simulated datasets as a function of decoder architecture. **(a)** Map-map FSC of final volumes generated from tomoDRGN homogeneous network training on simulated class E ribosomes with indicated decoder architectures against the corresponding ground truth volume. Panels correspond to different box and pixel sizes; colors correspond to different tomoDRGN model architectures. **(b)** Volumes were generated at each epoch during training and spatial frequencies at which map-map FSC=0.5 are plotted against cumulative wall time for models of different architectures (colors) on images of different box and pixel sizes (panels). Circles note total wall time elapsed and resolution achieved after 50 epochs of training.

**Extended Data Figure 4:**
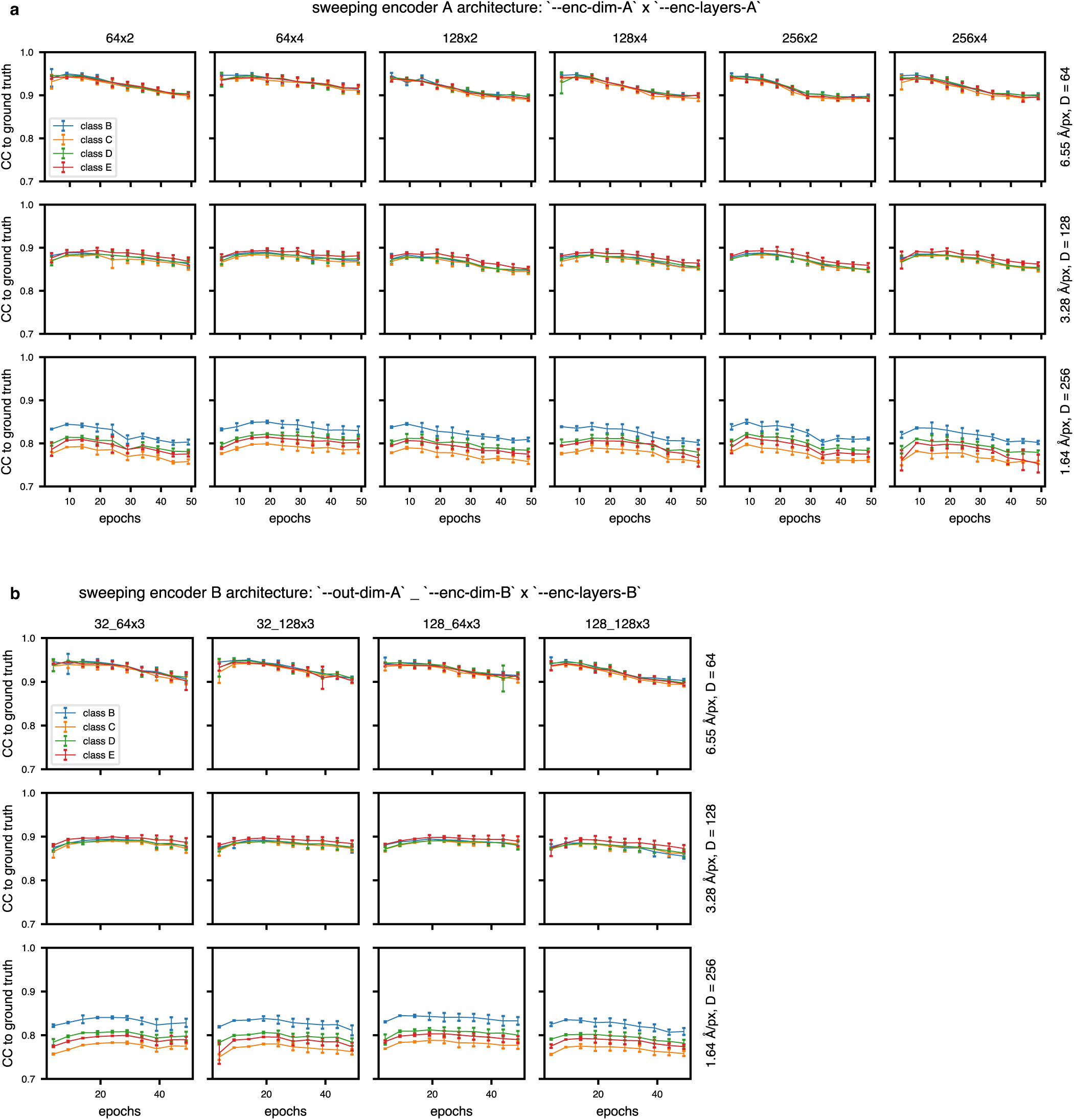
TomoDRGN model quality with heterogeneous simulated datasets as a function of encoder architectures. (a, b) Mean per-class volumetric correlation coefficient for tomoDRGN models trained with indicated encoder A architectures (panel A titles) or encoder B architectures (panel B titles). At 5 epoch intervals, 10 volumes from each volume class were generated and used to calculate volumetric correlation coefficients to the corresponding ground truth ribosome assembly intermediate volume. Error bars denote standard error of the mean in correlation coefficient among the tomoDRGN volumes at that epoch and the corresponding ground truth volume.

**Extended Data Figure 5:**
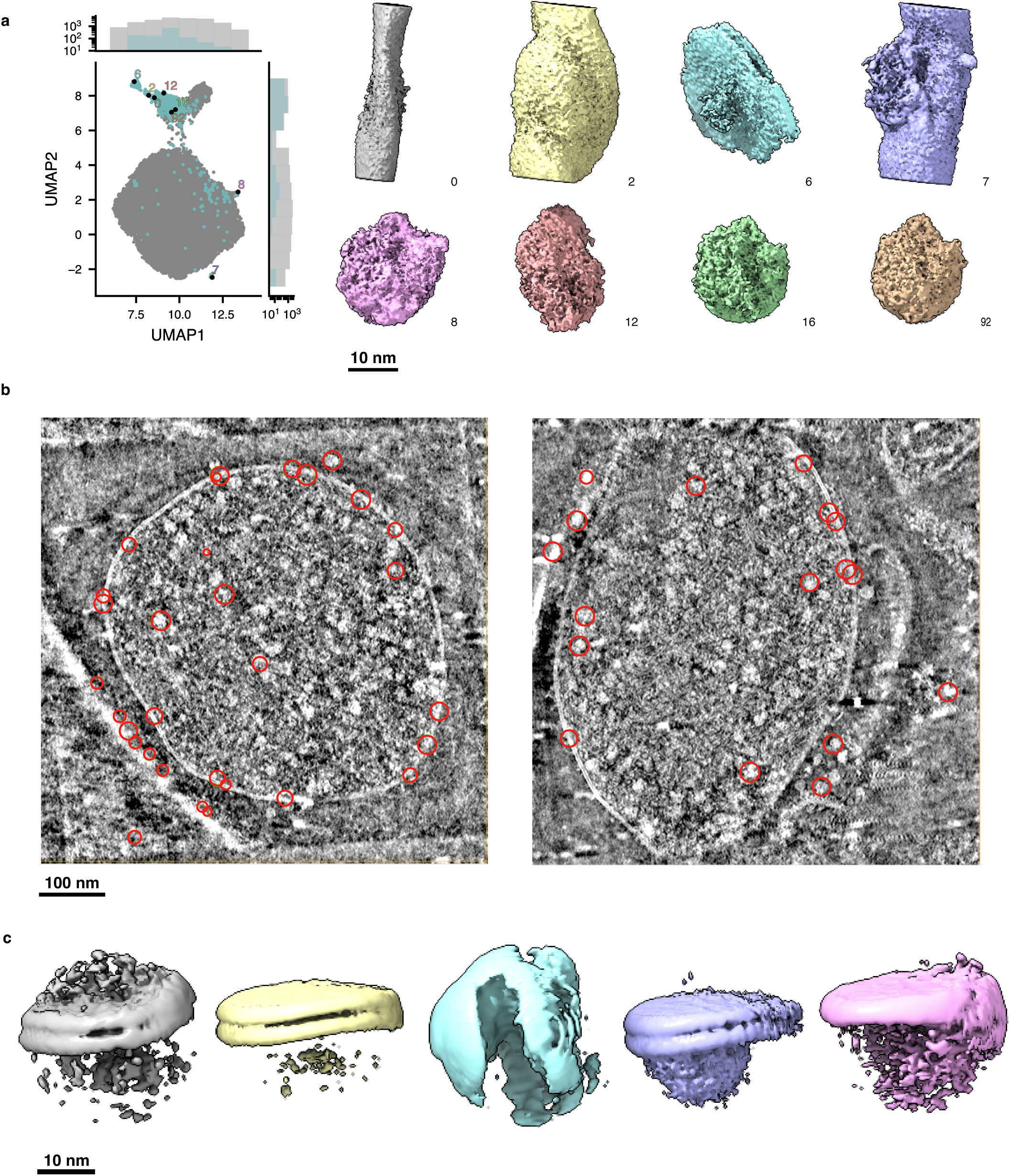
Using tomoDRGN to identify non-ribosomal particles picked from EMPIAR-10499 tomograms. **(a)** UMAP and corresponding sampled volumes from tomoDRGN heterogeneous network training from Fig. 4a. Eight representative non-ribosomal particles identified through manual inspection of *k*=100 *k*-means clustering of latent space are rendered at a constant isosurface and pose. **(b)** Two tomograms are shown in slice view using Cube (https://github.com/dtegunov/cube) with locations of particles labeled as non-ribosomal annotated within each tomogram. **(c)** RELION3-based multiclass (*k*=5) *ab initio* sub-tomogram volume generation using particles annotated as non-ribosomal (n=1,310).

**Extended Data Figure 6:**
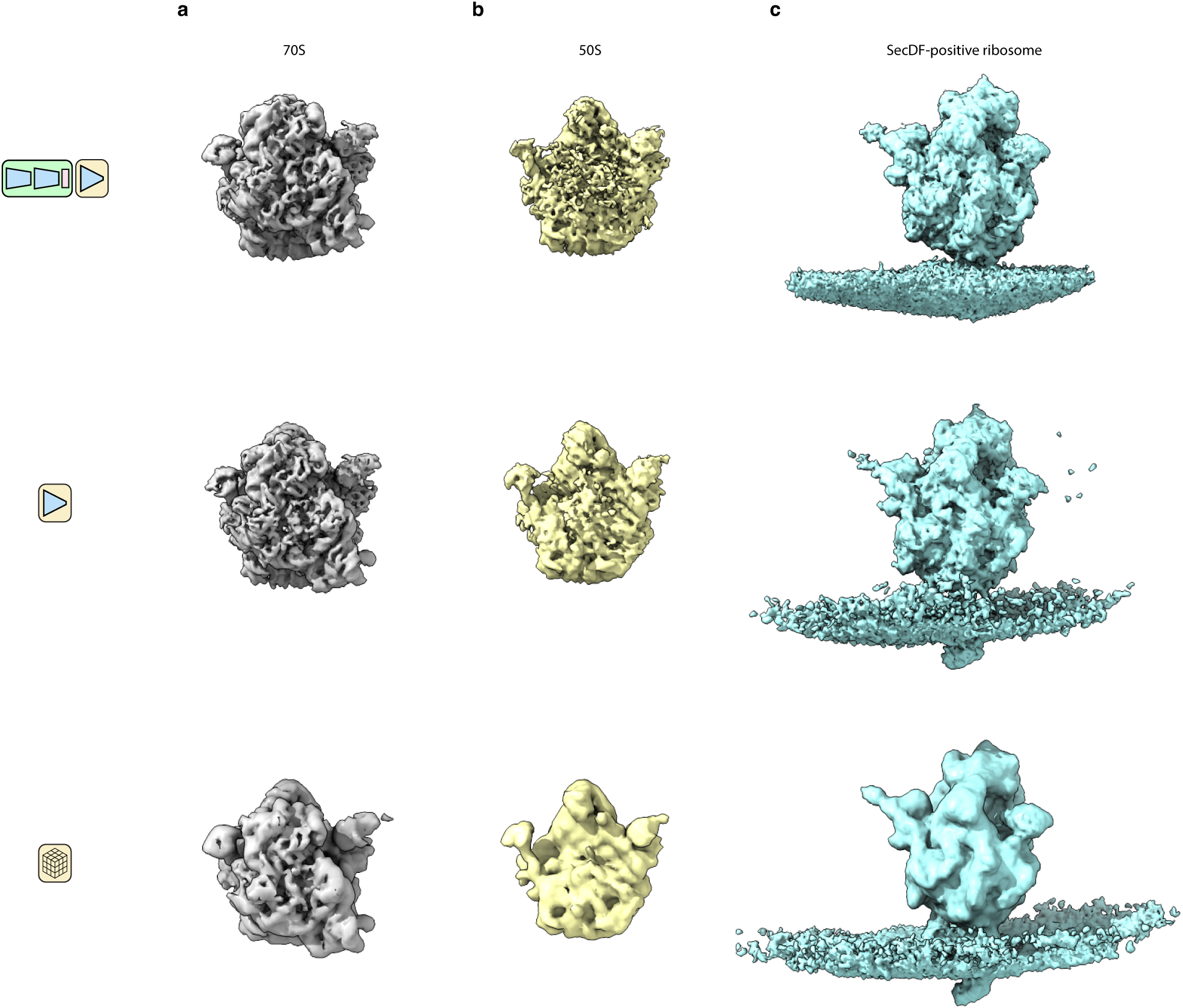
Validation of tomoDRGN-generated volumes. Comparison of volumes generated by a full tomoDRGN network (row 1), an isolated decoder neural network (row 2), or traditional sub-tomogram averaging (row 3). A full tomoDRGN network was trained on the heterogeneous ribosomal particle stack (row 1, n=20,981, see Figs. 4d and 5a) and representative volumes are depicted. Separate tomoDRGN homogeneous decoder networks were trained on one of three homogeneous substacks corresponding to **(a)** 70S particles (n=20,129); **(b)** 50S particles (n=852); or **(c)** SecDF-positive ribosomes (n=380). Traditional STA was also performed on each of these three particles stacks.

**Extended Data Figure 7:**
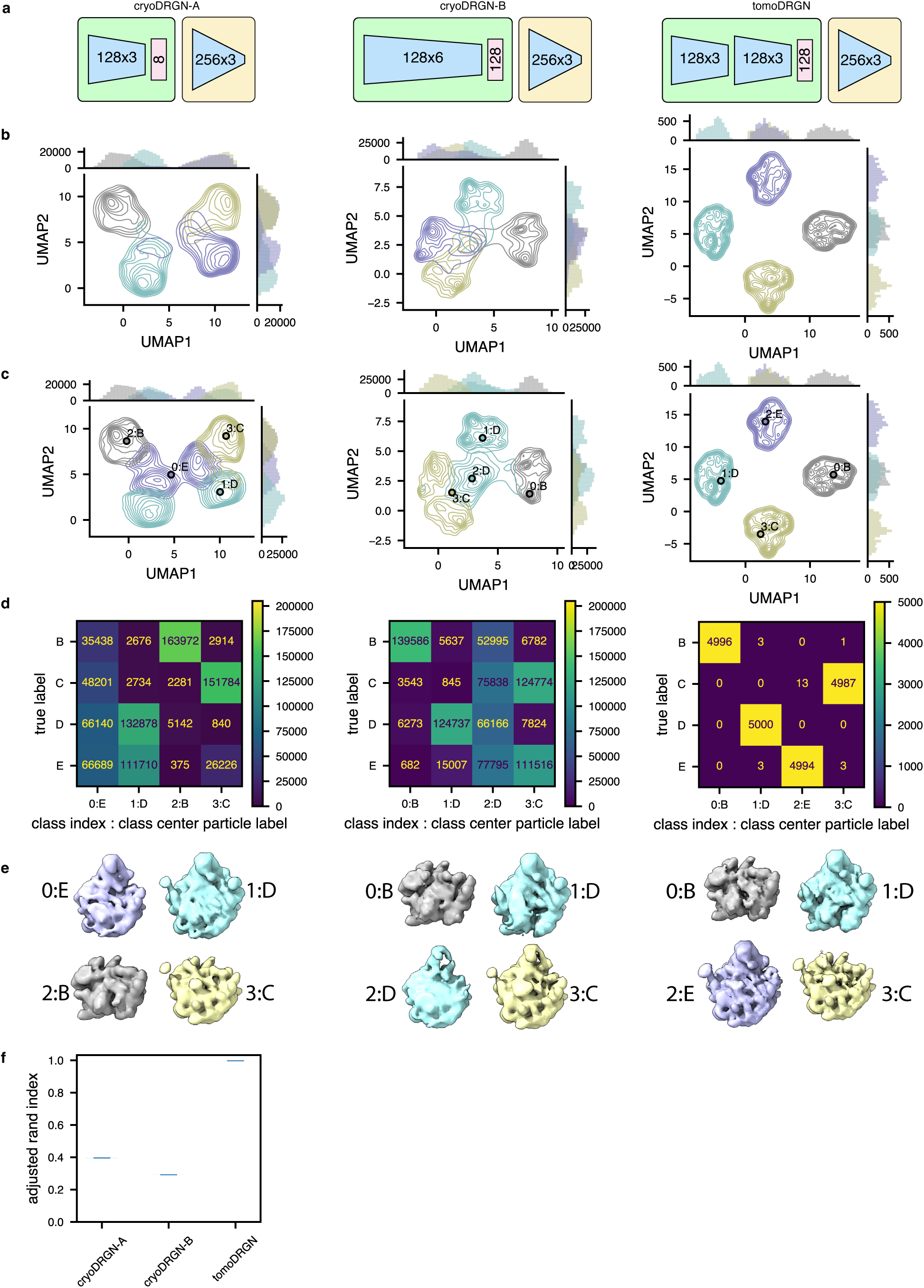
CryoDRGN fails to consistently encode structural heterogeneity using a simulated tilt series dataset. **(a)** Schematic of two cryoDRGN network architectures that were tested, and the tomoDRGN architecture used in Fig. 2c-e. Each model was trained using the same simulated dataset of ribosome large subunit assembly classes B-E (Davis *et al*., 2016) consisting of 41 tilt images for each of 5,000 particles for each of the four assembly states and thus the dataset was treated by cryoDRGN as n=820,000 images (see Methods). **(b)** UMAP of final epoch latent embeddings of each particle image, represented as a kernel density estimate (KDE) is plotted, with KDEs independently estimated and plotted for each of the four ground truth assembly states (bottom). **(c)** UMAP of final epoch latent embedding with *k*=4 *k*-means latent classification of the resulting latent space. KDEs were independently estimated and plotted for each of the four *k*-means classes. The predicted labels are annotated by both the *k*-means class index (0-3) and corresponding ground truth class label (B-E) of the central particle within each *k*-means class. **(d)** Confusion matrix of ground truth class labels versus *k*=4 *k*-means latent classification. **(e)** Volumes sampled at the *k*=4 *k*-means cluster centers illustrated in (c). Volumes are annotated by the *k*-means class index and ground truth class label and colored by the ground truth class label. **(f)** Violin plot of consistency of *k*=4 *k*-means clustering of each model by Adjusted Rand Index (Hubert and Arabie, 1985) (n = 100 randomly seeded initializations, higher values correspond to greater fidelity to ground truth classification).

**Extended Data Figure 8:**
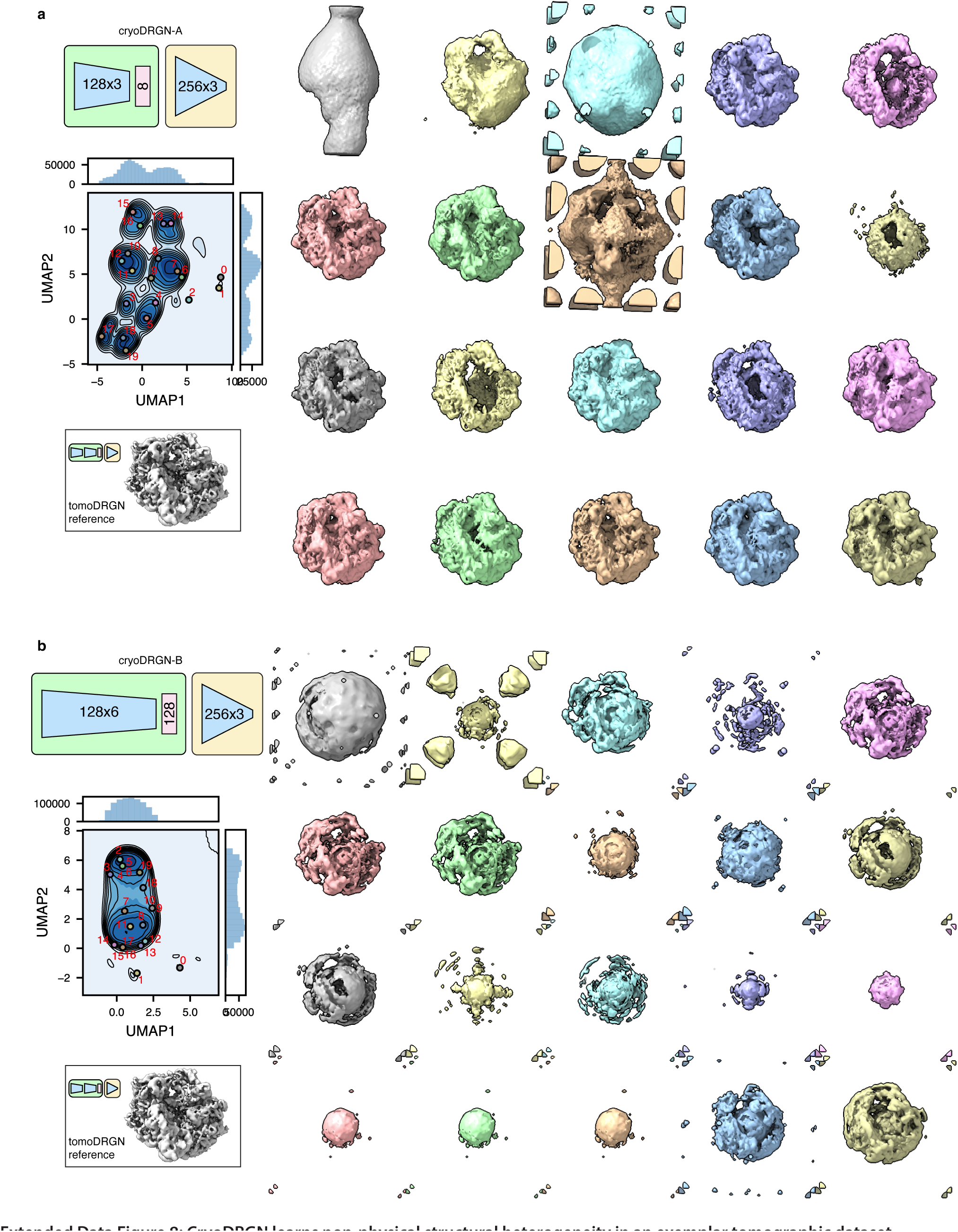
CryoDRGN learns non-physical structural heterogeneity in an exemplar tomographic dataset. Two cryoDRGN models **(a, b)** were trained on the unfiltered particle stack of *Mycoplasma pneumoniae* ribosomes from Fig. 4a (n = 22,291 particles, treated as n = 913,931 images). The latent space is shown as a KDE plot following UMAP dimensionality reduction, with *k*=20 *k*-means class center particles annotated (left) and corresponding volumes visualized (right). Note that many putative 70S particles lack density in the particle core. A reference 70S volume sampled from tomoDRGN’s model in Fig. 4a is shown in the same pose for comparison.

**Extended Data Figure 9:**
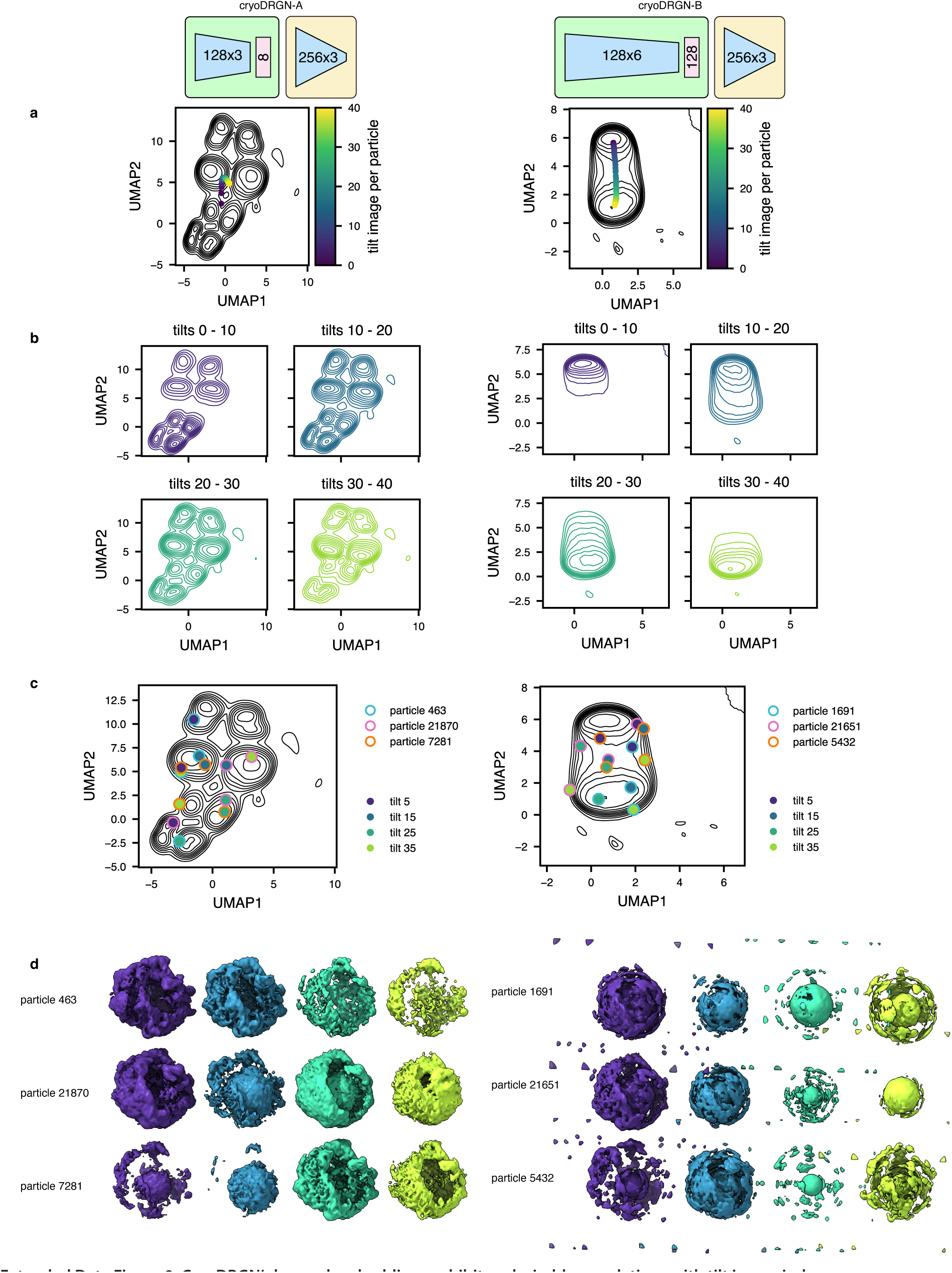
CryoDRGN’s learned embeddings exhibit undesirable correlations with tilt image index. **(a)** Two cryoDRGN models were tested on the unfiltered particle stack of *Mycoplasma pneumoniae* ribosomes from Fig. 4a. The latent space is shown as a KDE plot following UMAP dimensionality reduction. The latent embeddings were binned by the tilt image index, and the median value across each bin is annotated. **(b)** KDEs from panel A replotted after binning by tilt image index quartiles. **(c)** KDEs from panel A with annotated positions corresponding to three representative particles evaluated using their 5^th^, 15^th^, 25^th^, or 35^th^ tilt images. **(d)** Volumes generated from cryoDRGN using the latent embeddings highlighted in panel C.

**Table 1:** Impact of weighting and masking on performance of tomoDRGN homogeneous reconstruction. Summary statistics for tomoDRGN homogeneous network training using the simulated ribosome class E particles at different box and pixel sizes, in the presence or absence of reconstruction weighting and masking.

| Box size (px) | Pixel size (Å) | Weighting | Masking | Architecture | # Trainable parameters | # Data points per particle | Training time per 1k particles (min) | VRAM per particle (GB) | Max resolution (1/px) | Epochs to max resolution | Wall clock to max resolution (min) |
| --- | --- | --- | --- | --- | --- | --- | --- | --- | --- | --- | --- |
| 64 | 6.55 | - | - | 256 x 3 | 247,298 | 131,528 | 0.26 | 1.95 | 0.48 | 1 | 1.30 |
| 64 | 6.55 | + | - | 256 x 3 | 247,298 | 131,528 | 0.26 | 1.95 | 0.48 | 1 | 1.30 |
| 64 | 6.55 | - | + | 256 x 3 | 247,298 | 96,776 | 0.22 | 1.76 | 0.48 | 1 | 1.11 |
| 64 | 6.55 | + | + | 256 x 3 | 247,298 | 96,776 | 0.22 | 1.76 | 0.48 | 1 | 1.10 |
| 128 | 3.28 | - | - | 256 x 3 | 296,450 | 526,932 | 0.88 | 4.59 | 0.49 | 16 | 70.58 |
| 128 | 3.28 | + | - | 256 x 3 | 296,450 | 526,932 | 0.88 | 4.59 | 0.49 | 3 | 13.17 |
| 128 | 3.28 | - | + | 256 x 3 | 296,450 | 194,064 | 0.43 | 2.47 | 0.49 | 3 | 6.38 |
| 128 | 3.28 | + | + | 256 x 3 | 296,450 | 194,064 | 0.43 | 2.47 | 0.49 | 3 | 6.39 |
| 256 | 1.63 | - | - | 256 x 3 | 394,754 | 2,108,712 | 4.42 | 18.88 | 0.25 | 30 | 662.49 |
| 256 | 1.63 | + | - | 256 x 3 | 394,754 | 2,108,712 | 4.47 | 18.88 | 0.33 | 36 | 804.64 |
| 256 | 1.63 | - | + | 256 x 3 | 394,754 | 378,516 | 1.21 | 4.42 | 0.33 | 42 | 253.25 |
| 256 | 1.63 | + | + | 256 x 3 | 394,754 | 378,516 | 1.20 | 4.42 | 0.32 | 39 | 234.58 |

**Table 2:** Impact of network architecture on tomoDRGN homogeneous network reconstruction. Summary statistics for tomoDRGN homogeneous network training using the simulated ribosome class E particles at various box and pixel sizes, sweeping the number of nodes per layer in the decoder network.

| Box size (px) | Architecture | # Trainable parameters | # Data points per particle | Training time per 1k particles (min) | VRAM per particle (GB) | Max resolution (1/px) | Epochs to max resolution | Wall clock to max resolution (min) |
| --- | --- | --- | --- | --- | --- | --- | --- | --- |
| 64 | 64 x 3 | 24,962 | 96,776 | 0.15 | 1.42 | 0.48 | 1 | 0.75 |
| 64 | 128 x 3 | 74,498 | 96,776 | 0.17 | 1.54 | 0.48 | 1 | 0.85 |
| 64 | 256 x 3 | 247,298 | 96,776 | 0.22 | 1.76 | 0.48 | 1 | 1.11 |
| 64 | 512 x 3 | 887,810 | 96,776 | 0.37 | 2.15 | 0.48 | 1 | 1.84 |
| 64 | 768 x 3 | 1,921,538 | 96,776 | 0.58 | 2.79 | 0.48 | 1 | 2.88 |
| 128 | 64 x 3 | 37,250 | 194,064 | 0.28 | 1.85 | 0.38 | 43 | 60.74 |
| 128 | 128 x 3 | 99,074 | 194,064 | 0.33 | 2.04 | 0.49 | 15 | 24.94 |
| 128 | 256 x 3 | 296,450 | 194,064 | 0.43 | 2.47 | 0.49 | 4 | 8.69 |
| 128 | 512 x 3 | 986,114 | 194,064 | 0.75 | 3.33 | 0.49 | 2 | 7.48 |
| 128 | 768 x 3 | 2,068,994 | 194,064 | 1.22 | 4.53 | 0.49 | 1 | 6.12 |
| 256 | 64 x 3 | 61,826 | 378,516 | 0.88 | 3.48 | 0.20 | 42 | 185.81 |
| 256 | 128 x 3 | 148,226 | 378,516 | 0.94 | 3.67 | 0.26 | 47 | 221.67 |
| 256 | 256 x 3 | 394,754 | 378,516 | 1.15 | 4.42 | 0.32 | 33 | 190.30 |
| 256 | 512 x 3 | 1,182,722 | 378,516 | 1.81 | 6.10 | 0.34 | 22 | 199.42 |
| 256 | 768 x 3 | 2,363,906 | 378,516 | 3.61 | 11.92 | 0.35 | 20 | 360.84 |

**Table 3:**
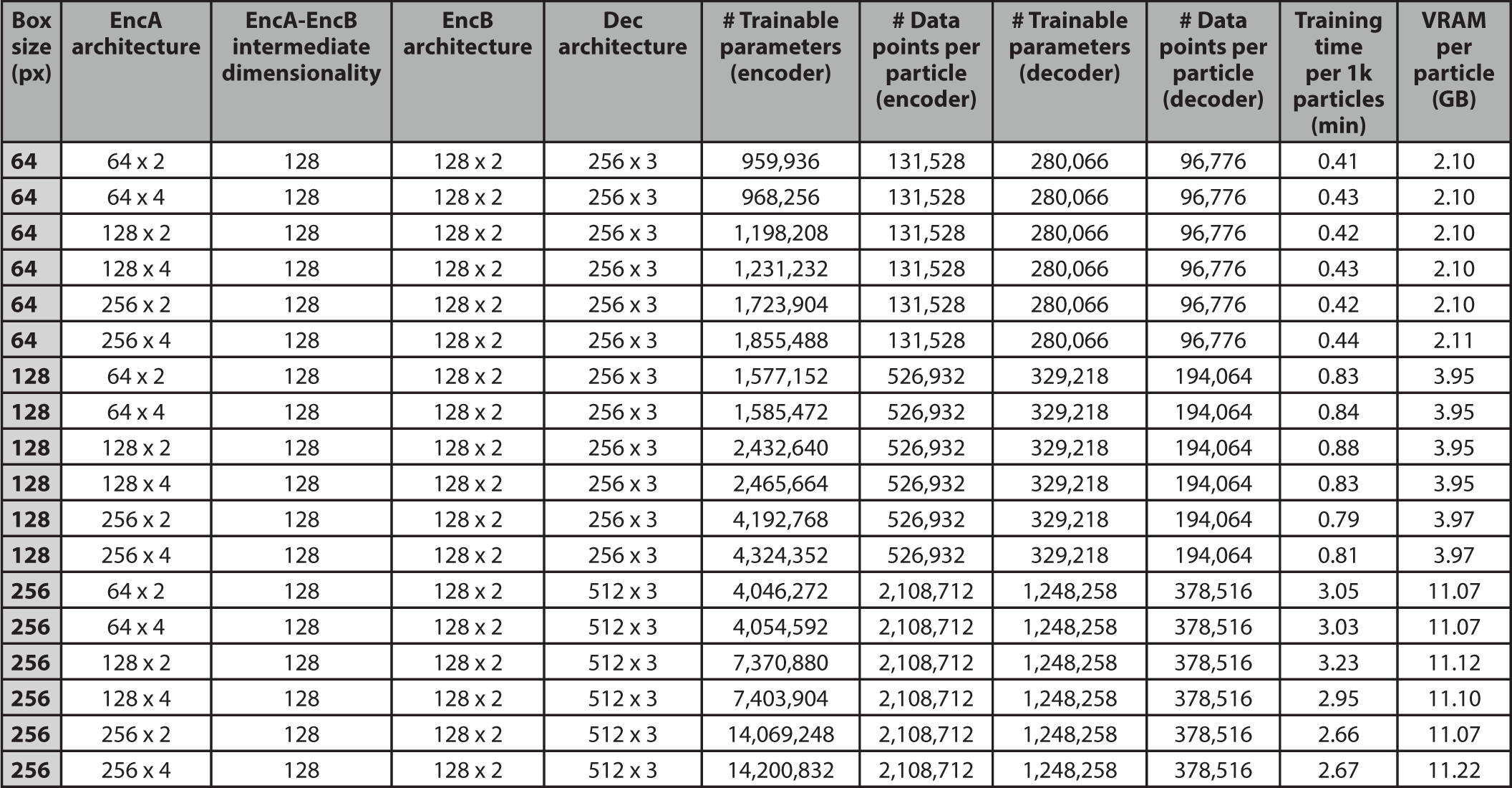
Impact of encoder network A architecture on tomoDRGN heterogeneous network reconstruction. Summary statistics for tomoDRGN heterogeneous network training using the simulated ribosome 4-class particles at various box and pixel sizes, sweeping the encoder A architecture (number of nodes per layer and number of layers).

**Table 4:** Impact of encoder network B architecture on tomoDRGN heterogeneous network reconstruction. Summary statistics for tomoDRGN heterogeneous network training using the simulated ribosome 4-class particles at various box and pixel sizes, sweeping the encoder A output layer size and the encoder B architecture (number of nodes per layer and number of layers).

| Box size (px) | EncA architecture | EncA-EncB intermediate dimensionality | EncB architecture | Dec architecture | # Trainable parameters (encoder) | # Data points per particle (encoder) | # Trainable parameters (decoder) | # Data points per particle (decoder) | Training time per 1k particles (min) | VRAM per particle (GB) |
| --- | --- | --- | --- | --- | --- | --- | --- | --- | --- | --- |
| 64 | 128 x 3 | 32 | 64 x 3 | 256 x 3 | 577,568 | 131,528 | 280,066 | 96,776 | 0.41 | 2.10 |
| 64 | 128 x 3 | 32 | 128 x 3 | 256 x 3 | 715,040 | 131,528 | 280,066 | 96,776 | 0.41 | 2.11 |
| 64 | 128 x 3 | 128 | 64 x 3 | 256 x 3 | 841,856 | 131,528 | 280,066 | 96,776 | 0.41 | 2.10 |
| 64 | 128 x 3 | 128 | 128 x 3 | 256 x 3 | 1,231,232 | 131,528 | 280,066 | 96,776 | 0.41 | 2.10 |
| 128 | 128 x 3 | 32 | 64 x 3 | 256 x 3 | 1,812,000 | 526,932 | 329,218 | 194,064 | 0.79 | 3.95 |
| 128 | 128 x 3 | 32 | 128 x 3 | 256 x 3 | 1,949,472 | 526,932 | 329,218 | 194,064 | 0.85 | 3.96 |
| 128 | 128 x 3 | 128 | 64 x 3 | 256 x 3 | 2,076,288 | 526,932 | 329,218 | 194,064 | 0.87 | 3.95 |
| 128 | 128 x 3 | 128 | 128 x 3 | 256 x 3 | 2,465,664 | 526,932 | 329,218 | 194,064 | 0.86 | 3.95 |
| 256 | 128 x 3 | 32 | 64 x 3 | 256 x 3 | 6,750,240 | 2,108,712 | 427,522 | 378,516 | 2.23 | 9.43 |
| 256 | 128 x 3 | 32 | 128 x 3 | 256 x 3 | 6,887,712 | 2,108,712 | 427,522 | 378,516 | 2.22 | 9.43 |
| 256 | 128 x 3 | 128 | 64 x 3 | 256 x 3 | 7,014,528 | 2,108,712 | 427,522 | 378,516 | 2.18 | 9.43 |
| 256 | 128 x 3 | 128 | 128 x 3 | 256 x 3 | 7,403,904 | 2,108,712 | 427,522 | 378,516 | 2.16 | 9.43 |

## Notes

### Competing Interest Statement

The authors have declared no competing interest.

